# Filament formation is a conserved mechanism of Thoeris SIR2 effector activation

**DOI:** 10.64898/2026.03.01.708072

**Authors:** Biswa P. Mishra, Veronika Masic, Tamim Mosaiab, Harshika Belani, Trent Conroy, Lou Brillault, Crystall M. D. Swarbrick, Zhiyu Zang, Bostjan Kobe, Yun Shi, Mark von Itzstein, Joseph P. Gerdt, Thomas Ve

**Affiliations:** Institute for Biomedicine and Glycomics, Griffith University, Gold Coast, QLD 4222, Australia; Department of Chemistry, Indiana University, Bloomington, IN 47405, USA; Centre for Microscopy and Microanalysis, The University of Queensland, Brisbane, QLD 4072, Australia; Biosecurity Research Program and Training Centre, Gulbali Institute, Charles Sturt University, Wagga Wagga, NSW 2678, Australia; School of Chemistry and Molecular Biosciences, Institute for Molecular Bioscience and Australian Infectious Diseases Research Centre, University of Queensland, QLD 4072, Australia; Global Health Institute, Swiss Federal Institute of Technology Lausanne (EPFL), Lausanne 1015, Switzerland; Medicinal Chemistry, Monash Institute of Pharmaceutical Sciences, Monash University, Parkville, Victoria 3052, Australia

## Abstract

Thoeris defence systems protect bacteria from phages via abortive infection. In type I Thoeris systems, ThsA effectors containing silent information regulator 2 (SIR2) and SMF/DprA-LOG (SLOG) domains are activated by the cyclic ADP-ribose (ADPR) isomer 3′cADPR, triggering abortive infection via nicotinamide adenine dinucleotide (NAD^+^) depletion. 3′cADPR activates the NADase activity of *Bacillus subtilis* ThsA tetramers via filament formation of its SIR2 domains, but the molecular details of how 3′cADPR triggers this process remain incompletely understood. Here, we demonstrate that ThsA activation by 3′cADPR-induced SIR2 filament formation is conserved in type I Thoeris systems from *Streptococcus equi* and *Entercoccus facium.* We present cryo-electron microscopy structures of the *S. equ*i ThsA filament bound to 3′cADPR and the non-cleavable NAD^+^ analog carba-NAD^+^, and of the *S. equi* ThsA tetramer bound to 3′cADPR. These structures reveal that SIR2 filament formation is required to stabilise an active site conformation that can bind and hydrolyse NAD^+^. The structures also show that 3′cADPR induces quaternary alterations in the SLOG dimers and consequently the SIR2 tetramer to enable ThsA filament formation. Collectively, our study provides a comprehensive understanding of 3′cADPR-induced activation of type I Thoeris effectors.

## Introduction

To combat phage infections, bacteria employ multiple defence strategies, including the use of nucleotide-based second messengers that trigger effector proteins and inhibit viral propagation (*1–4*). Thoeris is a family of defence systems that utilise Toll/interleukin-1 receptor (TIR) domains and nicotinamide adenine dinucleotide (NAD^+^) to trigger abortive infection (Abi) upon phage detection (*5, 6*). They consist of a single Abi-triggering effector (ThsA) gene and one or multiple TIR domain encoding (ThsB) genes (*5*) and occur in at least 8% of bacterial genomes.

Three different ThsA effector variants have been characterised in detail: a silent information regulator (SIR2) and SMF/DprA-LOG (SLOG) domain-containing effector (ThsA^SIR2-SLOG^; type I Thoeris); a transmembrane (TM) and macro domain-containing effector protein (ThsA^TM-macro^; type II Thoeris); and a caspase-like protease effector (type IV Thoeris) (*5, 7*). The ThsB genes encode TIR domain-containing proteins responsible for phage detection (e.g. by sensing capsid proteins) and production of signalling molecules that activate the ThsA effectors (*6, 8*). ThsB TIR domains have NAD^+^-cleavage activity and can produce a variety of signalling molecules such as 1′ ′-3′ cyclic ADP-ribose (3′cADPR), histidine-ADPR and N7-cADPR, which activate type I, II and IV Thoeris effectors, respectively (*7, 9–12*). Type II Thoeris effectors trigger abortive infection via membrane disruption, while Type IV Thoeris effectors halt phage infection by executing caspase-mediated cell-death. By contrast, type I Thoeris systems abort phage infection via rapid depletion of cellular NAD^+^, and 3′cADPR binding to ThsA^SIR2-SLOG^ effectors activates the NADase function of their SIR2 domains (*6, 9, 10, 13*). Recent reports have expanded the Thoeris family to include types III and V–XI, each associated with unique effector variants, some of which are activated by 2′cADPR and N1-cADPR (*14, 15*).

Biochemical studies of type I Thoeris systems from *B. cereus* MSX-D12, *Streptococcus equi* and *Enterococcus facium* revealed that their ThsA^SIR2-SLOG^ effectors exist as tetramers and that 3′cADPR binds to a conserved pocket in their SLOG domains (*10, 16*). Recently, Tamulaitiene et al demonstrated that the *B. cereus* ThsA (BcThsA) tetramers stack together via their SIR2 domains and form filaments in the presence of the activator 3′cADPR. Filament formation activates the NADase function of the SIR2 domain, leading to rapid NAD^+^ depletion and Abi (*13*). However, the mechanisms by which 3′cADPR promotes filament assembly, enables NAD^+^ access to the active site, and whether filament-mediated activation is conserved across other type I Thoeris systems, remains incompletely understood.

In this study, we show that a small proportion of *S. equi* ThsA (SeThsA) and *E. facium* ThsA (EfThsA) tetramers form filaments when incubated with 3′cADPR, and that these filaments have significantly higher NADase activity, when compared to the 3′cADPR-bound tetramers. A 2.7 Å resolution cryoEM structure of the SeThsA filament in complex with both 3′cADPR and the non-cleavable NAD^+^ analog, carba-NAD^+^, reveals stacked SeThsA tetramers in a similar arrangement to the BcThsA filament. Like BcThsA and our previously reported structure of inactive SeThsA, the tetrameric unit of the SeThsA filament has a core consisting of four SIR2 domains flanked by SLOG dimers at both sides. The carba-NAD^+^ binding mode is analogous to other SIR2 domains, suggesting a similar mechanism of NAD^+^ cleavage. Structural comparisons with the inactive, ligand-free SeThsA tetramer show that 3′cADPR binding induces changes to the SLOG dimer interface, which consequently lead to long-range conformational changes in the SIR2 tetramer interfaces. These quaternary rearrangements enable filament formation, resulting in active site conformational changes to permit NAD^+^ access and cleavage. We also report a 3.5 Å resolution cryoEM structure of the 3′cADPR-bound SeThsA tetramer. The structure demonstrates that 3′cADPR can also induce a different type of SIR2 tetramer rearrangement that does not enable filament formation and only affords sub-optimal binding of NAD^+^ to the active site, explaining the differences in NADase activity between the filament and tetramer assemblies. Collectively, our study demonstrates that 3′cADPR-induced higher-order oligomerisation is a conserved mechanism of activation for Thoeris type 1 effectors.

## Results

### SeThsA and EfThsA form filaments upon 3′cADPR binding

Tamulaitiene et al recently demonstrated that tetramers of the SIR2-SLOG ThsA effector from *Bacillus cereus* MSX-D12 (BcThsA) stack together, forming filaments in the presence of the activator 3′cADPR (*13*). Formation of these higher-order assemblies stabilises the active conformation of the BcThsA SIR2 domain, enabling rapid NAD^+^ depletion and phage protection.

To explore if higher-order assembly formation is a conserved mechanism of SIR2-SLOG ThsA effector activation, we analysed the oligomeric state of *E. faecium* ThsA (EfThsA; 45% sequence identity with BcThsA) and *S. equi* ThsA (SeThsA; 43% sequence identity with BcThsA) by size-exclusion chromatography. We have previously shown that ligand-free SeThsA and EfThsA exist as tetramers in solution (*10*). In the absence of 3′cADPR, these tetramers of SeThsA and EfThsA eluted at ∼11.6 mL on an S200 10/300 column (Fig. 1a and S1). After the addition of 3′cADPR to SeThsA and EfThsA, we observed new peaks at the void volume of the S200 column (Fig. 1a and S1). Negative-stain electron microscopy analysis revealed that these void-volume peaks of both SeThsA and EfThsA are composed of short filaments with a diameter of ∼20 nm and a length of 70-700 nm, while the major peaks at ∼11.5-11.6 mL consist of tetrameric particles (Fig. 1b). NMR-based NADase assays demonstrated that both the filament and tetramer fractions of SeThsA and EfThsA can cleave NAD^+^ into nicotinamide and ADPR, but the filament fractions have much higher NADase activity (Fig. 1c). Taken together, these findings support the model that ThsA activation by 3′cADPR involves higher-order oligomerisation.

**Fig. 1.**
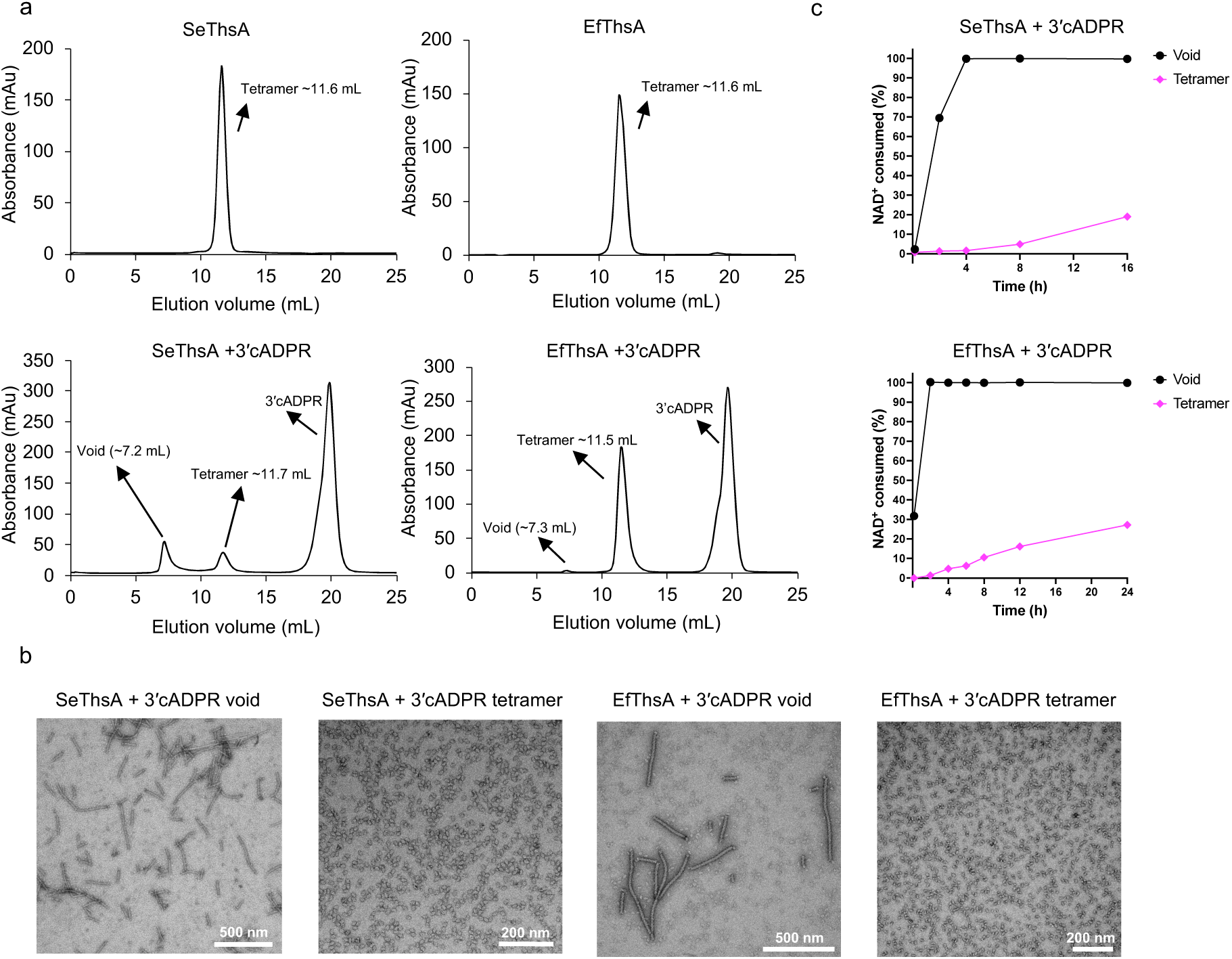
SeThsA and EfTshA filament formation and NADase activation by 3′cADPR. (a) Size-exclusion chromatography profiles of SeThsA +/- 3′cADPR and EfThsA +/- 3′cADPR. (b) Representative negative-stain EM micrographs of SeThsA and EfThsA void-volume and tetramer peaks. (c) NADase activities of SeThsA (top) and EfThsA (bottom) void-volume and tetramer peaks. The SeThsA and EfThsA concentrations were 0.2 μM and 0.1 μM, respectively. The initial NAD^+^ concentration was 500 μM. The experiments in (a) and (c) were conducted two times with similar results.

### CryoEM structure of the SeThsA filament

To provide structural insight into ThsA activation, we determined a cryoEM structure of SeThsA filaments incubated with the NAD^+^ analog carba-NAD^+^ at 2.82 Å resolution (Fig. 2a-c and S2). Carba-NAD^+^ is a non-cleavable analog of NAD^+^, in which a cyclopentane ring replaces the furanose of the nicotinamide-ribonucleotide moiety (*17, 18*). The reconstruction revealed stacked SeThsA tetramers in a helical arrangement with a twist of 120.22° and a helical rise of 43.63 Å (Fig. 2c-d). Similar to the crystal structure of the inactive ligand-free SeThsA tetramer (*10*) and the active BcThsA filament (*13*), the tetrameric unit of the SeThsA filament has D2 symmetry, with a core consisting of two SIR2 dimers flanked by SLOG dimers at both sides (Fig. 2e). Only the SIR2 domains are involved in intrafilament interactions; the SLOG domains form no helical contacts.

**Fig. 2.**
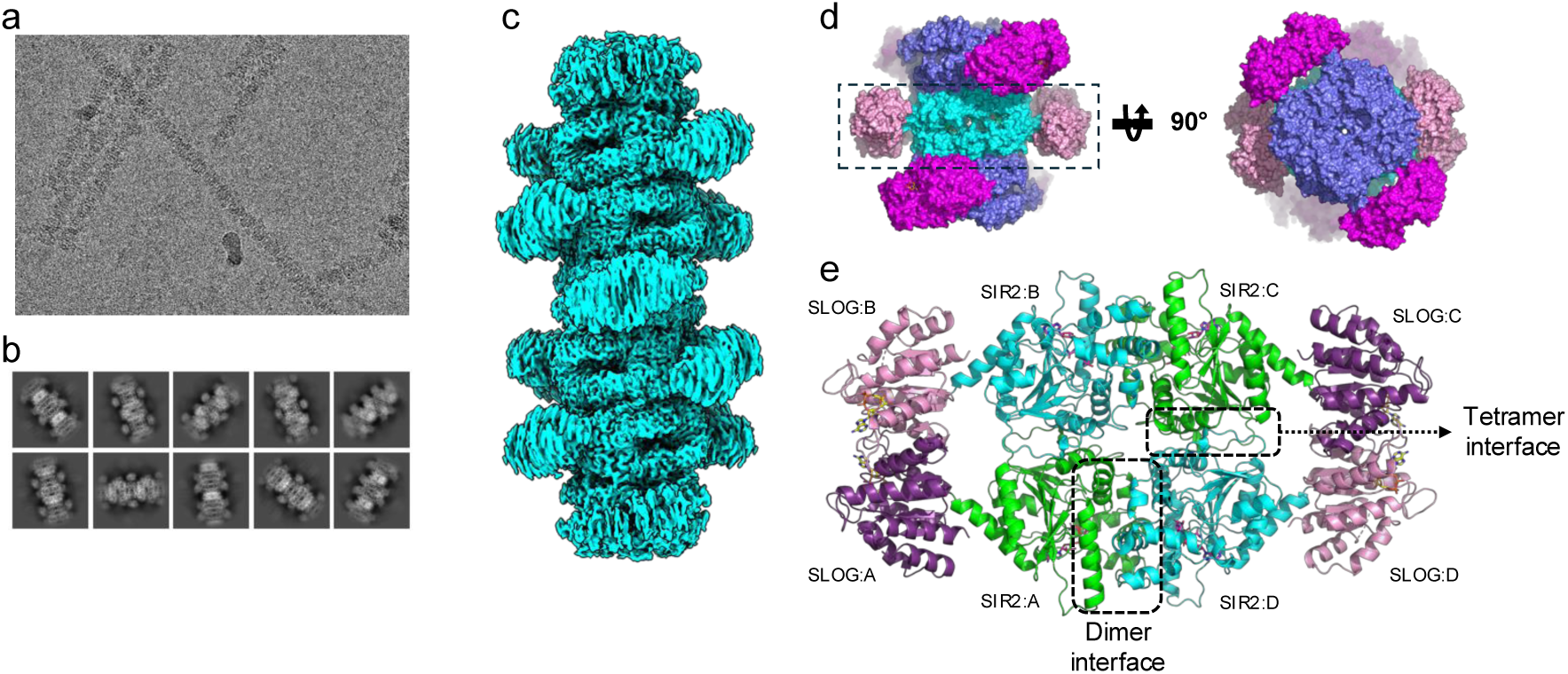
CryoEM structure of the SeThsA filament. (a) Representative micrograph. (b) Representative 2D classes. (c) Electrostatic potential density map of SeThsA filament. (d) Model of the SeThsA filament in surface representation. Three stacked tetramers are shown. The central tetramer is coloured in cyan (SIR2 domains) and pink (SLOG domains), while the top and bottom tetramers are coloured in slate (SIR2 domains) and magenta (SLOG domains). (e) Cartoon representation of an individual tetramer in the SeThsA filament. Carba-NAD^+^ and 3′cADPR (sticks) are highlighted in magenta and yellow, respectively.

### Mechanism of NAD^+^ binding

The SeThsA filament cryoEM map had well resolved densities for carba-NAD^+^ in the active site of the SIR2 domains (Fig. S2f). Similar to other SIR2 enzymes (*19, 20*), the substrate binds to a cleft located at the interface between the Rossmann fold and a small three-helix bundle domain, which is inserted in between the α2_1_ and α5 helices (Fig. 3a-b). The adenine group of the NAD^+^ mimic stacks against the side-chain of Y265, while the pyrophosphate group form hydrogen bonds with the side-and main-chain of S199 and the main-chain of A32. The nicotinamide riboside moiety is located in a deep pocket of the SIR2 domain formed by residues A32, I36, F41, S42, W43, F88, T112, N113, Y114, D115, H153, I166 and Y169. The NH_2_ group of the nicotinamide forms hydrogen bonds with the side-chain of D115 and the main-chain of Y114, while the ribose interacts with the side-chain of H153 and the main-chain of T112. The carba-NAD^+^ binding mode is analogous to the yeast Hst2:carba-NAD^+^ complex (*21*) (Fig. S3a), suggesting a similar mechanism of NAD^+^ cleavage involving formation of an oxocarbenium intermediate stabilised by the conserved N113 residue that is essential for ThsA NADase activity (*13, 16*).

**Fig. 3.**
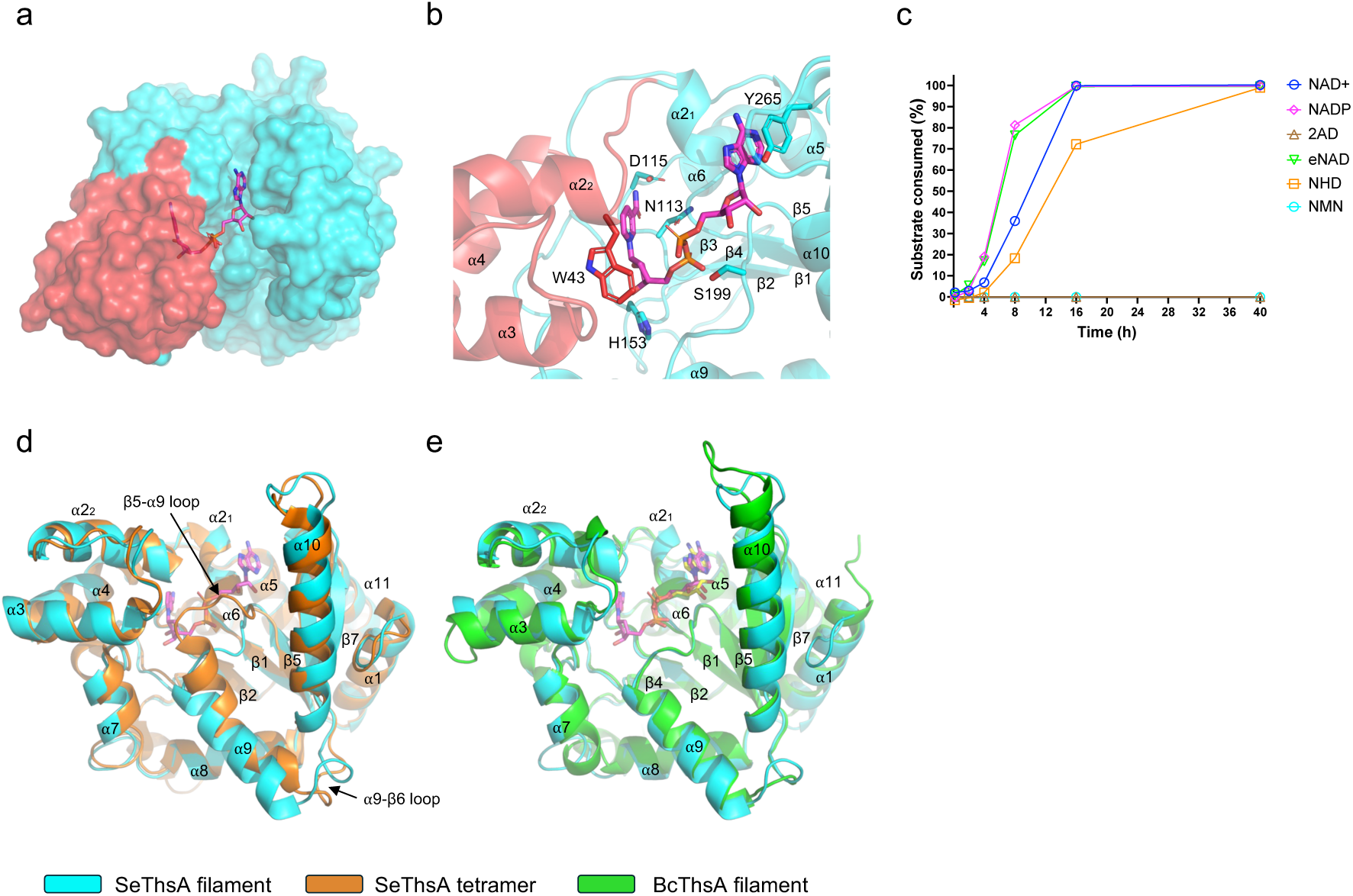
SeThsA active site. (a) Surface representation of the SIR2 domain in the SeThsA filament. The Rossman and three-helix domains are coloured in cyan and red, respectively. Carba-NAD^+^ is shown in stick representation (magenta). (b) Enlarged cutaway view of the SIR2 active site. (c) Reaction progress curves of 0.1 μM SeThsA + 500 μM substrate in the presence of 1 μM 3′cADPR. The experiments were conducted two times with similar results. (d-e) Structural superpositions of the SIR2 domain in the SeThsA filament with (d) the ligand-free inactive SeThsA tetramer (PDB: 7UXT) and (e) the BcThsA filament (PDB: 8BTO).

In the SIR2:carba-NAD^+^ complex, both the distal adenine base and adenine-linked ribose are exposed to solvent, while the nicotinamide ribose moiety is almost completely buried. It is thus possible that ThsA SIR2-SLOG effectors can also cleave dinucleotides with different distal bases or distal ribose modifications, but are less likely to tolerate modifications to the nicotinamide ribose moiety. Consistent with this hypothesis, SeThsA can efficiently cleave nicotinamide adenine dinucleotide phosphate (NADP^+^), etheno-NAD^+^ and nicotinamide hypoxanthine dinucleotide (NHD^+^), but not **2AD**, which has a 1,2-dihydro-2,7-naphthyridin-1-one base instead of nicotinamide (Fig. 3c). SeThsA cannot cleave nicotinamide mononucleotide, suggesting that the diphosphate group, adenine group and adenine-linked ribose are essential for substrate binding, which is consistent with our structural data.

Superposition of the SIR2 domain in the filament and ligand-free tetramer SeThsA structures (rmsd =1.4 Å for 275 Cα atoms) revealed a substantial shift (>5 Å) in the position of the β5-α9 loop and α9 helix (Fig. 3d). The side-chain of W43 in the active site region has also flipped in the filament structure to accommodate carba-NAD^+^. In the filament, the β5-α9 loop is an important component of the catalytic site, as it interacts with the pyrophosphate moiety of carba-NAD^+^. However, in the ligand-free tetramer structure, this loop is shifted upwards and carba-NAD^+^ cannot bind to the active site, due to steric clashes of its nicotinamide ribose and pyrophosphate group with the β5-α9 loop (Fig. 3d). The BcThsA filament structure also has similar β5-α9 loop and α9 helix conformations (Fig. 3e). By contrast, the α2_1_-α2_2_ loop, which has been proposed to control access to NAD^+^ in BcThsA (*13, 16*), has an almost identical open conformation in the tetramer and filament structures of SeThsA (Fig. S3b).

### 3′cADPR binding to the SLOG domains

The local resolution of the SLOG domain dimers was lower than that of the core SIR2 tetramer in our reconstruction (Fig. S2), suggesting that the linker region connecting the SLOG and SIR2 domains has some conformational flexibility. Symmetry expansion and local refinement of a masked SLOG dimer increased the resolution to 2.89 Å and the map had well resolved cryoEM density for 3′cADPR (Fig. S2). The overall structure of the SLOG domains is similar to both inactive ligand-free SeThsA (rmsd = 1.2 Å for 184 Cα atoms) and the BcThsA filament (rmsd = 1.8 Å for 180 Cα atoms) (Fig. 4a-b), and the 3′cADPR binding mode is almost identical between SeThsA and BcThsA (Fig. 4c).

**Fig. 4.**
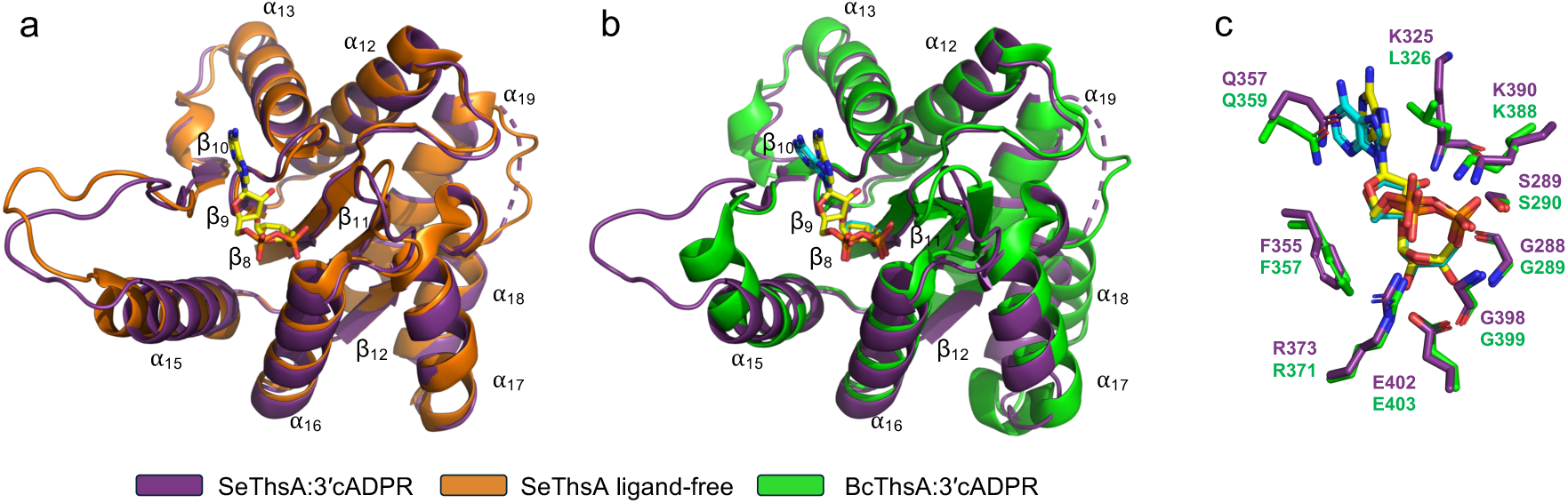
3′cADPR binding to the SLOG domain. (a-b) Structural superpositions of the SLOG domain in the SeThsA filament with (a) the ligand-free inactive SeThsA tetramer (PDB: 7UXT) and (b) the BcThsA filament (PDB: 8BTO). 3′cADPR is shown in yellow (SeThsA) and cyan (BcThsA) stick representation. (c) Comparison of the 3′cADPR binding site in the SeThsA (dark purple) and BcThsA (green) filaments.

### Structural alterations accompanying 3′cADPR binding

Comparisons of the filament and inactive ligand-free SeThsA structures reveal notable differences in both the SLOG dimers and SIR2 tetramer (Fig. 5a). The SLOG dimer interfaces in the SeThsA and BcThsA filaments are similar, but notably different to the ligand-free SeThsA tetramer (Fig. 5b). In both the SeThsA tetramer and filament structures, the SLOG dimer interface consists of residues in the α12-α14 helices, the β10 strand and the α13-β10 and β10-α14 loops (Fig. 5b). 3′cADPR binding induces conformational changes in the β10- α14 loop and α14 helix (Fig. 4a), and the two SLOG domains undergo both translation and rotational motion, bringing the two 3′cADPR molecules into closer proximity (Fig. 5b, Movie S1).

**Fig. 5.**
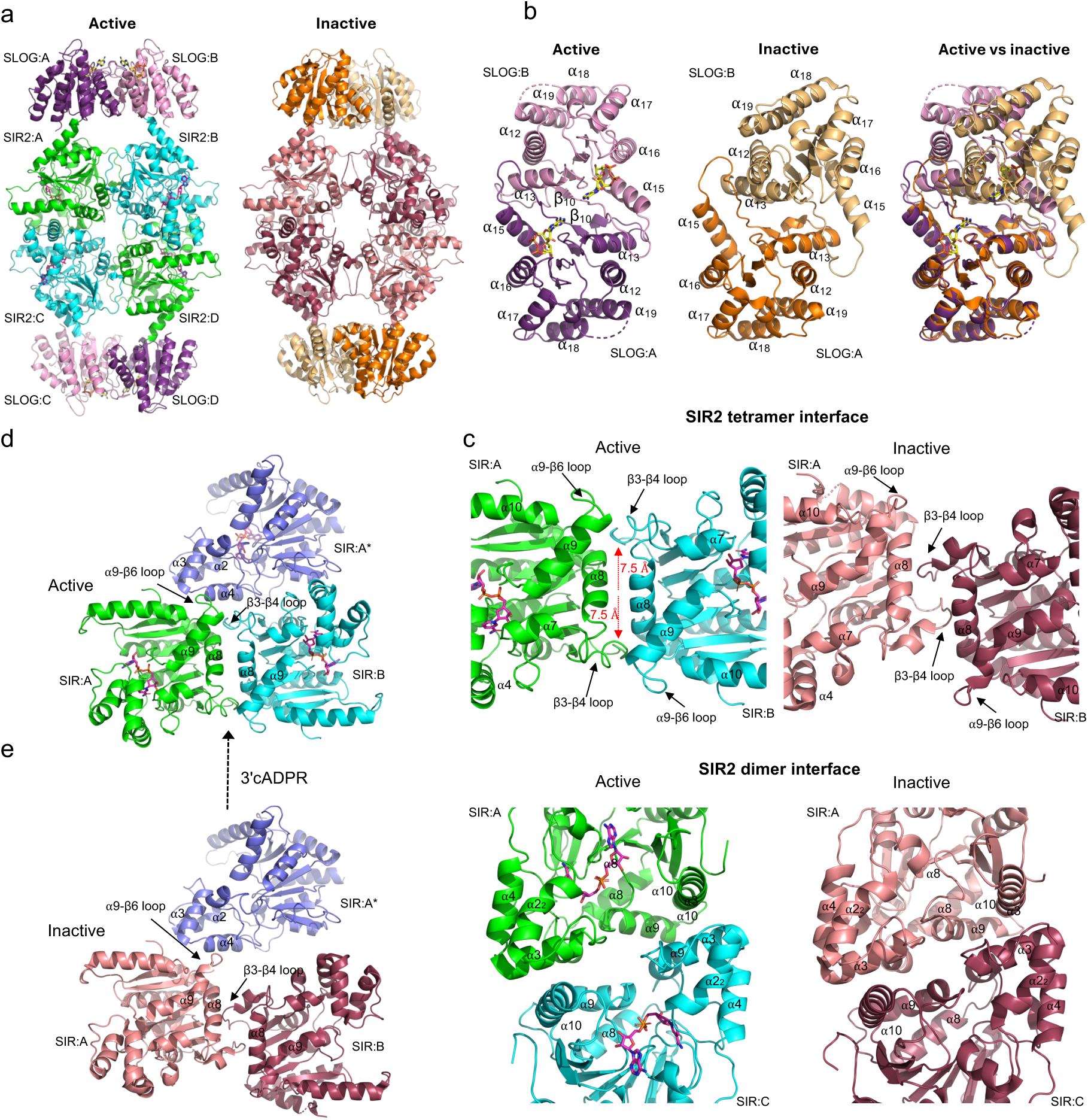
Changes in the SeThsA quaternary structure upon activation. (a) Structures of active (filament) and inactive (PDB: 7UXT) SeThsA tetramers. Carba-NAD^+^ and 3′cADPR (sticks) are highlighted in magenta and yellow, respectively. In each tetramer, the two SLOG dimers are identical and the SIR2 A:B and A:C interfaces are identical to the SIR2 C:D and B:D interfaces, respectively. (b) Comparison of SLOG dimers in the active (filament) and inactive (PDB: 7UXT) SeThsA tetramers. The SLOG dimers were superimposed using SLOG:A. (c) Comparison of the SIR2 tetramers in the active (filament) and inactive (PDB: 7UXT) SeThsA tetramers. Top panels: SIR2 tetramer interface. Bottom panels: SIR2 dimer interface. (d) Helical interface in the SeThsA filament. The SIR2 domain from the adjoining tetramer (SIR2:A*) is coloured in slate. (e) Hypothetical model of the helical interaction in (d) using the inactive (PDB: 7UXT) SeThsA tetramer. The model shows that 3′cADPR induce quaternary rearrangements that are required to create a stable SIR2 helical interface.

The 3′cADPR-induced alterations in the SLOG dimer result in major changes to the SIR2 A:B and C:D tetramer interfaces (Fig. 5c, Movie S2). In the ligand-free inactive SeThsA tetramer, these symmetric interfaces, with a buried surface area of ∼1244 Å^2^, consist of residues in the β3-β4 loops and α8 helices. Upon 3′cADPR-induced changes in the SLOG dimer, the SIR2 domains move ∼ 7.5 Å in opposite directions, which brings the α8 helices closer to each other and enables the β3-β4 loops to engage with residues at the tip of the α1 and α9 helices and the β5-α9 loops (Fig. 5c). The rearrangements also lead to an increased buried surface area of ∼2048 Å^2^. By contrast, the SIR2 A:D and B:C dimer interfaces only undergo minor rearrangements (Fig. 5c and Movie S3).

### SeThsA helical interface

The alterations to the SIR2 A:B and C:D interfaces create a new surface that enables SIR2 tetramers to stack together and form a filament (Fig. 5d). The β3-β4 and α9-β6 loops at the SIR2 A:B and C:D interfaces of each tetramer serve as docking surfaces for the three-helix bundle domain (α2-α4) and the β4-α7 loop of adjoining tetramers (Fig. 5d and 6a). The helical interface also involves residues in the α4-α5 loop, the β3 and β6 strands and the α1 and α10 helices, and has a buried surface area of 2809 Å^2^ (Fig. 6a). Although the BcThsA filament has an almost identical helical interface (Fig. S4a), many interacting residues are not conserved (Fig. S4b-c). Alanine substitution of two lysine residues in the SeThsA helical interface (K76 and K223) abolished 3′cADPR-induced filament formation and SeThsA activation (Fig. 6b and S5). The mutant also compromised activation of Thoeris antiviral defence in *Bacillus subtilis* (Fig. 6c). Cells expressing ThsB *from B. cereus* MSX-D12 (BcThsB) (*5*) together with the K76AK223A mutant of SeThsA produced ≥1000-fold more SPO1 plaques than those expressing BcThsB with wild-type SeThsA. Despite this, the K76AK223A mutant still conferred some protection, as plaque formation was ∼100-fold lower compared to cells expressing the catalytically inactive N113A SeThsA mutant (*16*) or cells lacking SeThsA entirely. We speculate that in *B. subtilis* cells with high ThsA expression, the K76AK223A mutant may still assemble into higher-order oligomers, thereby activating its NADase activity.

**Fig. 6.**
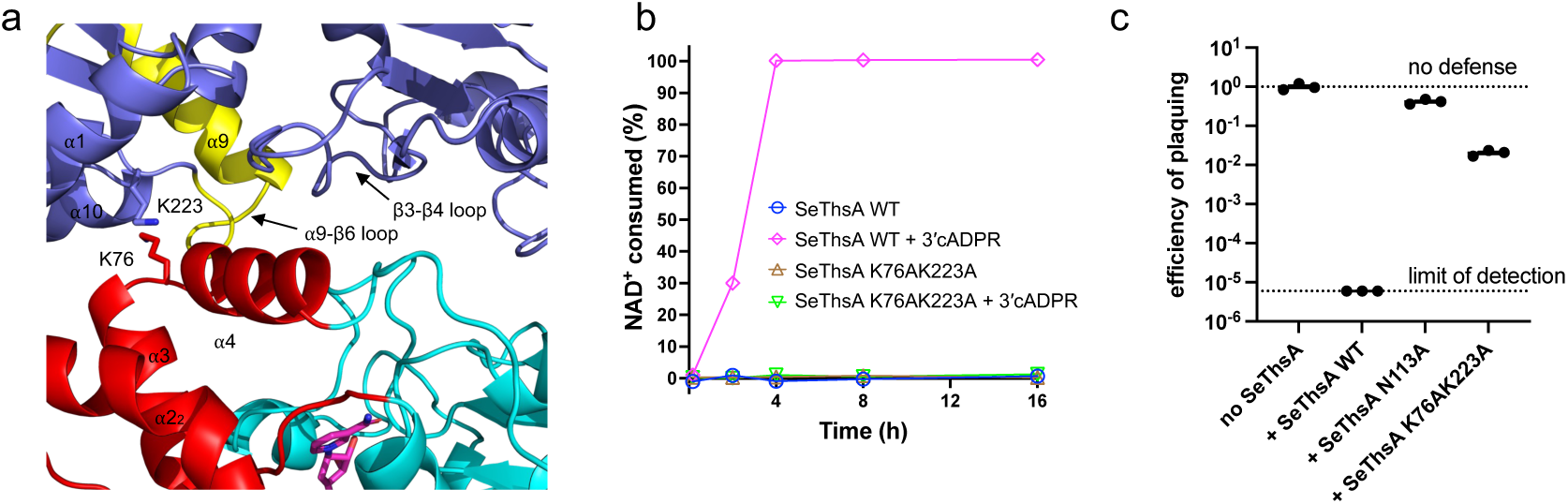
Filament formation is required for SeThsA activation. (a) Enlarged cutaway view of the SeThsA helical interface. The three-helix bundle domain is highlighted in red, while the SIR2 domains from the adjoining tetramer are displayed in slate. The α9 helix, and the β5-α9 and α9-β6-loops are highlighted in yellow. Mutated interface residues (K76 and K223) are shown as sticks. (b) NADase activities of 0.2 μM SeThsA +/- 1 μM 3′cADPR and SeThsA K76AK223A mutant +/- 1 μM 3′cADPR. The initial NAD^+^ concentration was 500 μM. The experiments were conducted two times with similar results. (c) Phage SPO1 challenge of *Bacillus subtilis* expressing BcThsB and SeThsA. Efficiency of plaquing was calculated relative to cells containing only BcThsB. No individual plaques were observed upon addition of wild-type SeThsA. Both mutations significantly improved plaquing efficiency, with the K76AK223A recovery being only partial. Mean ± SE of a biological triplicate is displayed, with each replicate designated by a circle.

Together, these results demonstrate that 3′cADPR-induced ThsA filament formation is a conserved mechanism of Thoeris defence activation.

### CryoEM structure of a SeThsA tetramer in complex with 3′cADPR

Analytical gel-filtration experiments showed that, after incubation with 3′cADPR, a large proportion of SeThsA and EfThsA elute as a tetramer (Fig. 1a-b). These tetrameric particles can still hydrolyse NAD^+^, but the activity is substantially lower compared to the SeThsA and EfThsA filaments (Fig. 1c). To provide insight into the molecular basis for these differences in NADase activity, we determined a 3.7 Å resolution cryoEM structure of a SeThsA tetramer after incubation with 3′cADPR (Fig. 7a and S6). The reconstruction revealed that 3′cADPR binding induces identical conformational changes in the SLOG domain dimers in the tetramer and the filament (Fig. 6b), but there are significant differences in the tertiary and quaternary structures of the SIR2 domains (Fig. 6c-d). The β5-α9 loop and α9 helix of the SIR2 domains in the 3′cADPR-bound tetramers have a similar conformation to the crystal structure of the inactive ligand-free SeThsA tetramer, suggesting that NAD^+^ cannot readily access the active sites (Fig. 6e). However, in two of the SIR2 subunits (chain B and C), the three-helix bundle domains are disordered, resulting in solvent exposure to the active site region that binds to the nicotinamide ribose portion of NAD^+^ (Fig. 6f). It is thus possible that NAD^+^ can bind sub-optimally to these active sites, explaining the low NADase activity observed for the 3′cADPR bound SeThsA tetramer (Fig. 1c).

**Fig. 7.**
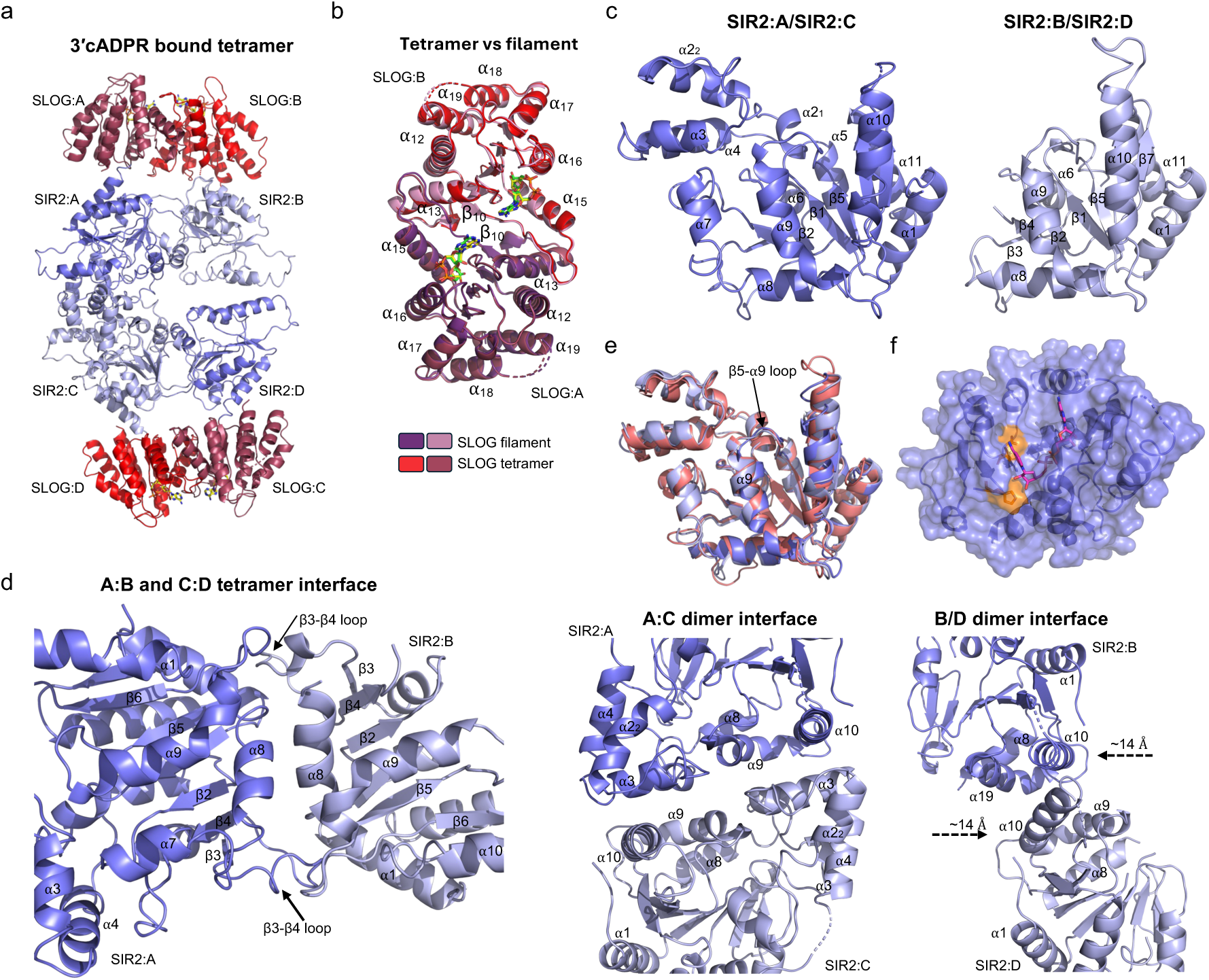
CryoEM structure of 3′cADPR-bound SeThsA tetramer. (a) Cartoon representation of the 3′cADPR bound tetramer. The SLOG domains are coloured red and raspberry; the SIR2 domains are coloured slate and light-blue. 3′cADPR (yellow) is shown in stick representation. The two SLOG dimers are identical. The SIR2 A:B and C:D tetramer interfaces are also identical, while the SIR2 A:C and B:D dimer interfaces have notably different configurations (see (f) for details). (b) Comparison of SLOG dimers in the SeThsA filament and 3′cADPR-bound tetramer. The SLOG dimers were superimposed using SLOG:A. (c) Comparison of the SIR2 domains in the 3′cADPR-bound tetramer. In SIR2:B and SIR2:D, the three-helix bundle domain (residues 92-98), the β4-α7 loop and α7 helix (residues 155-177) are disordered. (d) Structural superpositions of the SIR2 domain in the ligand-free inactive SeThsA tetramer (PDB: 7UXT) with the A (slate) and B (light-blue) chains of the 3′cADPR-bound SeThsA tetramer. (e) Surface representation of SIR2:B. The catalytically important N113 and H153 residues are highlighted in orange. (f) Quaternary structure of the SIR2 tetramer. Left panel: SIR2 A:B and C:D tetramer interfaces. Right panel: SIR2 A:C and B:D dimer interfaces. In the SIR2 B:D dimer interface, the two chains have moved in opposite directions, bringing the α10 helices into closer proximity.

Both the SIR2 A:B and C:D interfaces of the 3′cADPR-bound tetramer have undergone similar rearrangements as observed in the filament (Fig. 6d). The SIR2 A:C dimer interfaces are also similar in the 3′cADPR-bound tetramer and filament structures, but the SIR2 B:C dimer interfaces are significantly different (Fig. 6d) In the 3′cADPR-bound tetramer, the B and D SIR2 chains have moved in opposite directions, bringing the α10 helices into closer proximity (Fig. 6d). In this configuration, the three-helix bundle domains are no longer stabilised by α3-α10 interactions, resulting in their disordered conformation and consequently active site exposure. A similar configuration of SIR2 domains was observed in the crystal structure of BcThsA (Fig. S7) (*16*). These observations suggest that 3′cADPR can trigger two different types of SIR2 quaternary structure rearrangements in SeThsA.

## Discussion

In this study, we demonstrate that the NAD^+^ cleavage activity of ThsA effectors from *S. equi* and *E. facium* is activated by 3′cADPR-induced filament formation. We determined the cryoEM structures of SeThsA:3′cADPR complexes in both filamentous and tetrameric states, and by comparing these structures to the inactive, ligand-free SeThsA tetramer we reported previously (*10*), we found that 3′cADPR binding alters the SLOG dimer interfaces in SeThsA, which consequently induces long-range conformational changes to the core SIR2 tetramer. Our structures suggest that the SIR2 tetramer can be rearranged in two ways by 3′cADPR (Fig. 8):

**Fig. 8.**
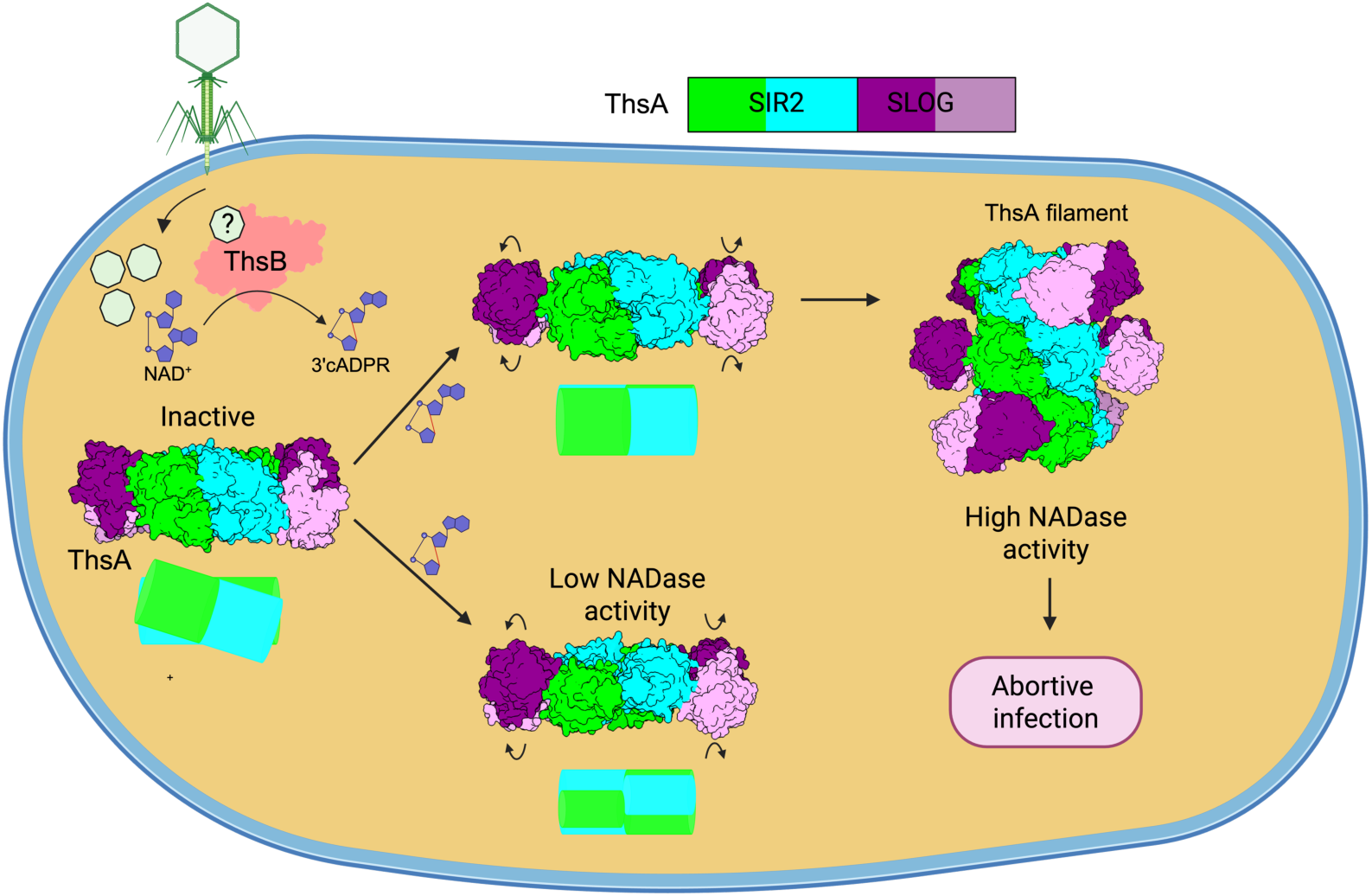
Model for the mechanism of action of Thoeris defense systems with a SIR2-SLOG ThsA effector. The figure was created with BioRender (www.biorender.com).

(i) the SIR2 tetramer interface is altered, leading to self-association of SeThsA tetramers into a filament and consequently a shift in the position of β5-α9 loops and α9 helices (Fig. 3d), which in turn enables NAD^+^ to enter the active sites; (ii) one of the dimer interfaces is also altered, preventing filament formation and resulting in destabilisation of the three-helix bundle domains, leading to solvent exposure of the catalytic N113 and H153 residues (Fig. 7f). Our NMR kinetics data (Fig. 1c) show that SeThsA and EfThsA filaments have high NADase activity. Alterations to the SeThsA helical interface abolished NADase activity and compromised antiphage defence, suggesting that filament formation via SIR2 tetramer interface alterations is required for Thoeris anti-phage activity, consistent with the BcThsA activation model proposed by Tamulatiene et al (*13*).

Based on structural comparisons of ligand-free SeThsA (PDB: 7UXT) and the crystal structure of BcThsA (PDB: 6LHX), we previously reported that 3′cADPR-induced activation of SeThsA likely involves SIR2 dimer interface alterations, destabilisation of the three-helix bundle domain and exposure of catalytic residues (*10*). Although our cryoEM data show that 3′cADPR can induce such changes to the SeThsA SIR2 tetramer, our kinetics data demonstrate that 3′cADPR bound SeThsA tetramers only have low NADase activity, which is unlikely to trigger NAD^+^ depletion and anti-phage defence.

Under our *in vitro* conditions, only a small population of 3′cADPR-bound SeThsA and EfThsA tetramers formed filaments and the major fraction of SeThsA and EfThsA remained as tetramers with low NADase activity. We speculate that cellular conditions may favour filament formation, but further studies will be required to visualise ThsA filament formation upon 3′cADPR binding in bacterial cells.

The structural basis for SIR2 NADase activation has also recently been reported for DSR2 (*22–26*), SIR2-HerA (*19, 27–29*), AVS (*30*), and SIR2-APAZ/Ago (*20, 31–33*) defence systems. Like Thoeris, these systems feature SIR2-domain oligomers in the active state. SIR2-APAZ/Ago SIR2 domains undergo a monomer-to-tetramer transition upon activation, while AVS SIR2 domains form filaments. DSR2 and SIR2-HerA SIR2 domains exist as preformed tetramers and dodecamers, respectively, that undergo small movements in the three-helix bundle domain upon activation. DSR2 and Thoeris have related tetramer architectures, whereas the repeating SIR2 dimer unit in the AVS filament is similar to the SIR2 dimer in Thoeris. The SIR2 interfaces in the SIR2-HerA dodecamer and the SIR2-APAZ/Ago tetramer are very different from Thoeris, DSR2 and AVS. Similar to Thoeris, the NAD^+^ cleavage site is located within one SIR2 domain in all of these systems, and substrate access and/or cleavage require conformational changes, which are regulated via SIR2 domain oligomerisation or rearrangements of preformed SIR2 oligomers. These SIR2 activation mechanisms are distinct to TIR NADases, the second major class of NAD^+^-depleting effectors involved in antiphage defence (*3, 34–39*), which have the active site located at the interface between two TIR domains (*10, 12*) and therefore require TIR self-association to form active sites. However, despite their differences, both systems are variations of the SCAF (signalling by cooperative assembly formation) mechanism (*40*).

In summary, our structural and biochemical data offer insights into the molecular mechanism of the Thoeris immune system, particularly regarding how 3′cADPR-induced alterations in the SLOG domains leads to SIR2 domain quaternary changes and substrate access to the active site.

## Materials and Methods

### Cloning

The SeThsA K76AK223A mutant DNA was made synthetically (gBlock, Integrated DNA Technologies) and cloned into the pMCSG7 vector using LIC (*41*). Pure plasmids were prepared using the QIAprep Spin Miniprep Kit (Qiagen) and the sequences confirmed by the Australian Genome Research Facility.

### Protein production and purification

SeThsA and SeThsA K76AK223A DNA, inserted into the pMCSG7 vector (N-terminal His6-tag, TEV (tobacco etch virus) protease cleavage site) were produced in *E. coli* BL21 (DE3) cells, using the autoinduction method (58) and purified to homogeneity, using a combination of immobilised metal-ion affinity chromatography (IMAC) and size-exclusion chromatography (SEC), as previously described (*10*). The cells were grown at 37°C, until OD600 of 0.6-0.8 was reached. The temperature was then reduced to 20°C, and the cells were grown overnight for approximately 16 h. The cells were harvested by centrifugation at 5000 x *g* at 4°C for 15 min and the cell pellets were resuspended in 2-3 mL of lysis buffer (50 mM HEPES pH 8.0, 500 mM NaCl) per g of cells. The resuspended cells were lysed using a digital sonicator and clarified by centrifugation (15,000 x *g* for 30 min). The clarified lysate was supplemented with imidazole (final concentration of 30 mM) and then applied to a nickel HisTrap column (Cytiva) pre-equilibrated with 10 CVs (column volumes) of the wash buffer (50 mM HEPES pH 8.0, 500 mM NaCl, 30 mM imidazole) at a rate of 4 mL/min. The column was washed with 10 CVs of the wash buffer, followed by elution of bound proteins using elution buffer (50 mM HEPES pH 8, 500 mM NaCl, 250 mM imidazole). The elution fractions were analysed by SDS-PAGE and the fractions containing the protein of interest were pooled and further purified on a S200 HiLoad 26/600 column pre-equilibrated with gel-filtration buffer (10 mM HEPES pH 7.5, 150 mM NaCl). The peak fractions were analysed by SDS-PAGE (sodium dodecyl sulfate–polyacrylamide gel electrophoresis), and the fractions containing SeThsA were pooled and concentrated to final concentrations of approximately 39.5 mg/mL (SeThsA), and 53 mg/mL (SeThsA K76AK223A), flash-frozen as 10 μL aliquots in liquid nitrogen, and stored at-80°C

### Analytical size-exclusion chromatography

For analysis of ThsA oligomerisation upon 3′cADPR binding, samples of SeThsA and EfThsA (25-100 μM) were incubated with 2 mM 3′cADPR for 1 h at room temperature in running buffer (10 mM HEPES pH 7.5, 150 mM NaCl). After incubation, 500 μL of each sample was loaded onto a S200 10/300 column (Cytiva) using an Akta Pure device (Cytiva) at 0.5 mL/min. Analysis of chromatograms and image preparation was performed in Unicorn 7 software (Cytiva) and Excel (Microsoft). Peak fractions were analysed by SDS-PAGE and negative-stain EM.

### Negative-stain EM

Four μL of sample was placed on a carbon-coated copper grid and incubated for 60 s. The grid was then washed with Milli-Q® H_2_O and stained with 1% uranyl acetate for 60 s and air-dried. The images were collected on a JEOL JEM-1400Flash TEM 120 kV transmission electron microscope at 20,000-60,000x magnification at 80 keV.

### CryoEM data collection

*SeThsA tetramer*: SeThsA protein (3.4 mg/mL) was incubated with 1 mM 3′cADPR for 1 h at room temperature and loaded onto a S200 26/60 SEC column (Cytiva) pre-equilibrated with gel-filtration buffer. Negative-stain EM analysis revealed that the right-hand shoulder of the major peak (Fig S6A) contained mostly tetramers (0.28 mg/mL) and was loaded onto a glow-discharged Quantifoil grid (R2/1 300-mesh holey carbon film coated, ProSciTech), blotted for 5.5 s under 95% humidity at 4°C, and plunged into liquid ethane, using a FEI Vitrobot Mark IV automatic plunge freezer (Thermo Fisher Scientific). For data collection, movies were acquired on a JEM-Z300FSC (Cryo-ARM300, JEOL Ltd.) operating at 300 keV equipped with an in-column Omega energy filter. Movies were recorded with a K3 Summit direct electron detector (Gatan) operating at 100,000x magnification (0.48 Å per pixel). All movies were exposed using a total dose of 40 e^-^/Å^2^ over 40 frames and a defocus range between-0.5 and - 2.5 µm.

*SeThsA filament*: SeThsA protein (1 mg/mL) was incubated with 1 mM 3′cADPR for 1 h at room temperature and loaded onto a Superdex 200 10/300 Increase SEC column (Cytiva) pre-equilibrated with 10 mM Hepes pH 7.5, 150 mM NaCl. The filament-containing peak at the void volume (Fig. S2a) was concentrated to 1.9 mg/mL and incubated with 1 mM carba-NAD^+^ for 1 h at 25°C. This sample was then loaded onto a glow-discharged Quantifoil grid (R1.2/1.3 300-mesh holey carbon film coated, ProSciTech), blotted for 5.5 s under 95% humidity at 4°C, and plunged into liquid ethane, using a FEI Vitrobot Mark IV automatic plunge freezer (Thermo Fisher Scientific). For data collection, movies were acquired on a JEM-Z300FSC (Cryo-ARM300, JEOL Ltd.) operating at 300 keV equipped with an in-column Omega energy filter. Movies were recorded with a K3 Summit direct electron detector (Gatan) operating at 100,000x magnification (0.48 Å per pixel). All movies were exposed using a total dose of 40 e^-^/Å^2^ over 40 frames and a defocus range between-0.5 and-2.5 µm.

### CryoEM data processing

All processing steps were performed using CryoSPARC v4.5.1 (*42*) and the cryoEM processing workflows are summarised in Figs. S2 and S6.

*SeThsA filament*: A total of 4,033 movies were collected and imported into CryoSPARC (*42*). Alignment of movie frames was performed using patch-based motion correction. Fitting of the contrast transfer function and defocus estimation was performed using patch-based CTF estimation. Initial 2D classes were generated from particles picked using the Blob picker and were used for template-based picking. A total of 3,241,637 particles were extracted with a 352-pixel box size. Several rounds of 2D classification were subsequently performed to remove inferior particles. After 2D classification, good particles were further classified into three 3D maps using ab initio reconstruction, followed by homogenous refinement and 3D classification (10 classes). The final set consisted of 238,570 particles and reached 2.82 Å resolution (after reference-based motion correction and local CTF refinement) using a helical refinement protocol with an initial volume from the best 3D class, a twist of 120.22°, a rise of 43.36 Å, and D2 symmetry. To perform local refinement of the SLOG dimer, the symmetry was expanded to C1, yielding 3,815,520 particles. A volume map for mask creation for local refinement was generated in UCSF ChimeraX using Segger (*43, 44*). A soft padded mask was generated using Volume Tools in CryoSPARC and was used for local refinement, yielding a 2.89 Å resolution SLOG dimer map.

*SeThsA tetramer*: A total of 2,408 movies were collected and imported into CryoSPARC. Alignment of movie frames was performed using patch-based motion correction. Fitting of the contrast transfer function and defocus estimation was performed using patch-based CTF estimation. Initial 2D classes were generated from particles picked using the Blob picker and were used for template-based picking. A total of 973,157 particles were extracted with a 256-pixel box size. Several rounds of 2D classification were subsequently performed to remove inferior particles. After 2D classification, good particles were further classified into three 3D maps using ab initio reconstruction followed by heterogenous refinement. The final set consisted of 131,258 particles and reached 3.71 Å resolution, using a non-uniform refinement protocol with C1 symmetry after reference-based motion correction. To perform local refinement, volume maps of the two SLOG dimers were generated in UCSF ChimeraX using Segger (*43, 44*). Soft-padded masks were generated using Volume Tools in CryoSPARC and were used for local refinement, yielding 3.71 Å and 3.65 Å maps of the two SLOG dimers respectively.

### Model building and refinement

*SeThsA filament*: The structure of the SeThsA tetramer was automatically built using ModelAngelo (*45*). The helical refinement map was used for building the SIR2 domains, while the local refinement map was used to build the SLOG domains. Following ModelAngelo, the SeThsA tetramer model was subjected to iterative rounds of model building and refinement using Coot (*46*) and phenix.real_space_refine from the PHENIX suite (*47, 48*). Ligand restraints for 3′cADPR and carba-NAD^+^ were generated using Grade2 (*49*) and the ligands were fitted manually in Coot, followed by refinement using phenix.real_space_refine. Two additional SeThsA tetramers were fitted into the helical refinement map using ChimeraX (*43*) and the final assembly was refined using phenix.real_space_refine (rigid body and ADP refinement). Model validation was performed using the phenix.validation_cryoem tool (*50*).

*SeThsA tetramer*: The SIR2 and SLOG domains from the filament structure were fitted into the tetramer reconstruction using ChimeraX (*43*). For the SIR2 domains, fitting was performed using the global map, while the SLOG domains were positioned using locally refined maps of the SLOG dimers. The resulting SIR2 domain tetramer underwent iterative model building and refinement using Coot (*46*) and the phenix.real_space_refine tool from the PHENIX suite (*47, 48*). In the final stage, the SLOG domain dimers were fitted into the global tetramer map, and the complete tetramer model was refined using phenix.real_space_refine, applying only rigid body and atomic displacement parameter (ADP) refinement. Model validation was performed using the phenix.validation_cryoem tool (*50*).

Statistical data for cryoEM data collection, refinement and validation are provided in Table S1. Structures were visualised and figures were generated using UCSF ChimeraX (*43*) and PyMol (Schrodinger).

### NMR-based enzymatic assays

NMR samples were prepared in 175 μL HBS buffer (50 mM HEPES, 150 mM NaCl, pH 7.5 in H_2_O), 20 μL D_2_O, and 5 μL DMSO-d6, resulting in a total volume of 200 μL. Each sample was subsequently transferred to a 3 mm Bruker NMR tube. All ^1^H NMR spectra were acquired with a Bruker Avance Neo 600 MHz NMR spectrometer equipped with a QCI-F cryoprobe at 298 K. To suppress resonance from H_2_O, a water suppression pulse program (P3919GP) using a 3-9-19 pulse-sequence with gradients (*51*) was implemented to acquire spectra with an acquisition delay of 2 s and 32 scans per sample. For each reaction, spectra were recorded at multiple time points such as 10 min, 2 h, 4 h, 8 h, and 16 h, depending on instrument availability. All spectra were processed by TopSpin™ (Bruker) and MNova (Mestrelab Research). The reaction progression was calculated based on the integration of non-overlapping resonance peaks, which vary depending on sample composition, from pairs of substrates and products, such as NAD^+^ and nicotinamide, respectively. The detection limit (signal-to-noise ratio > 2) was estimated to be 10 μM.

### Carba-NAD^+^ synthesis

Carba-NAD^+^ was synthesised according to a previously published method (*18*) and the ^1^H and ^13^C NMR spectra were in excellent agreement with previous reports (*17, 18*).

### Testing of SeThsA mutants in *B. subtilis*

*Construction of strains.* The BcThsB gene was cloned into pDR110 (Bacillus Genomic Stock Center #ECE311, see Table S2 for full plasmid sequence with insert) and transformed into *Bacillus subtilis* RM125 (Bacillus Genomic Stock Center #1A253), as described previously (*52*), selecting on LB + 100 µg/mL spectinomycin. The insertion was validated by PCR with primers 5′–ATGGCGAAAAGAGTTTTTTTTAGT–3′ and 5′–TTTAAAAGGAGAACCCACTTGATT–3′. The wild-type, N113A, and K76AK223A mutants of SeThsA were synthesized by Gene Universal and cloned into pMS039 (gift from Xindan Wang, see Table S2 for full plasmid sequences with inserts). All three plasmids were transformed into *B. subtilis* DK1042, as described previously (*53*), selecting on LB + chloramphenicol (5 µg/mL). Subsequently, each SeThsA-expressing region was transduced with phage SPP1 (*54*) into the recipient RM125:BcThsB strain, selecting on LB + chloramphenicol (5 µg/mL) supplemented with 10 mM sodium citrate. The inserts were PCR validated with primers 5′–GGAGATCGAAAAGGCAATAAA–3′ and 5′–GCCTGTATACTCAATAGGAAG–3′.

*Efficiency of plaquing.* The sensitivity of bacteria containing BcThsB only, BcThsB + SeThsA, BcThsB + SeThsA N113A, and BcThsB + SeThsA K76AK223A hosts to phage SPO1 on solid media was determined by the small drop plaque assay. In brief, 200 μL of an overnight culture of *B. subtilis* host was mixed with 5 mL 55 °C LB + 0.5% agar containing 1 mM IPTG and 0.5% xylose and poured on top of an LB + 1.5% agar plate. After an hour, 5 µL of 1:10 dilutions of SPO1 in LB was dropped on top of the soft agar. After the phage spot dried, the plate was incubated at 37 °C overnight. Plaques were counted to generate plaque-forming units (PFU) per mL of lysate for each host. Each PFU/mL measurement was normalized by the average PFU/mL value on the BcThsB-only host, where that host without SeThsA is defined as “efficiency of plaquing” = 1.

## Data availability

The cryo-EM density maps have been deposited in the Electron Microscopy Data Bank (EMDB) under accession numbers EMD-75058 [https://www.ebi.ac.uk/pdbe/entry/emdb/ EMD-75058] (SeThsA filament), EMD-75059 [https://www.ebi.ac.uk/pdbe/entry/emdb/EMD-75059] (SeThsA filament, local refinement map of SLOG dimer), EMD-75060 [https://www.ebi.ac.uk/pdbe/entry/emdb/ EMD-75060] (SeThsA tetramer), EMD-75061 [https://www.ebi.ac.uk/pdbe/entry/emdb/ EMD-75061] (SeThsA tetramer, local refinement map of SLOG dimer 1) and EMD-75062 [https://www.ebi.ac.uk/pdbe/entry/emdb/ EMD-75062] (SeThsA tetramer, local refinement map of SLOG dimer 2). The atomic coordinates have been deposited in the Protein Data Bank (PDB) under accession numbers 10CC [https://doi.org/10.2210/pdb 10CC/pdb] (SeThsA filament), 10CD [https://doi.org/10.2210/pdb 10CD/pdb] (SeThsA filament, local refinement map of SLOG dimer) and 10CE [https://doi.org/10.2210/pdb 10CE/pdb] (SeThsA tetramer). Protein structures used for analysis in this study are available in the Protein Data Bank under accession numbers 7UXT [https://doi.org/10.2210/pdb7UXT/pdb], 8BTO [https://doi.org/10.2210/pdb8BTO/pdb], 6LHX [https://doi.org/10.2210/pdb6LHX/pdb], and 1SZC [https://doi.org/10.2210/pdb1SZC/pdb].

## Acknowledgements

We acknowledge the Centre for Microscopy and Microanalysis, University of Queensland and staff. We thank Dr. Dan Kearns and Chengqian Zhang for assistance with *B. subtilis* cloning. The work was supported by the National Health and Medical Research Council (Investigator Grant 1196590 and 2042089 to TV; 2025931 to BK; Investigator Grant 1196520 and 2009677 to MvI), the Australian Research Council (Future Fellowship (FT200100572) to TV; Laureate Fellowship (FL180100109) to BK; and Discovery Early Career Research Award (DE250101258) to YS) and the National Science Foundation CAREER award (IOS-2143636 to JPG).

## Author Contributions

Writing – original draft, TV; Conceptualisation, TV; Investigation, BPM, VM, TM, HB, LB, CS, TC, ZZ, YS, TV; Writing – review & editing, all authors; Resources, TV, MvI; Methodology, BPM, VM, TM, HB, LB, CS, TC, ZZ, YS, TV; Data curation, YS, TM; Validation, BPM, VM, TM, HB, LB, CS, TC, ZZ, YS, TV; Funding acquisition, TV, JPG, MvI, YS, BK; Supervision, TV, JPG, MvI, YS, BK; Formal analysis, YS, TM; Visualisation, BPM, HB, YS, JPG, TV; Project administration, TV

## Supplementary Figures

**Fig. S1.**
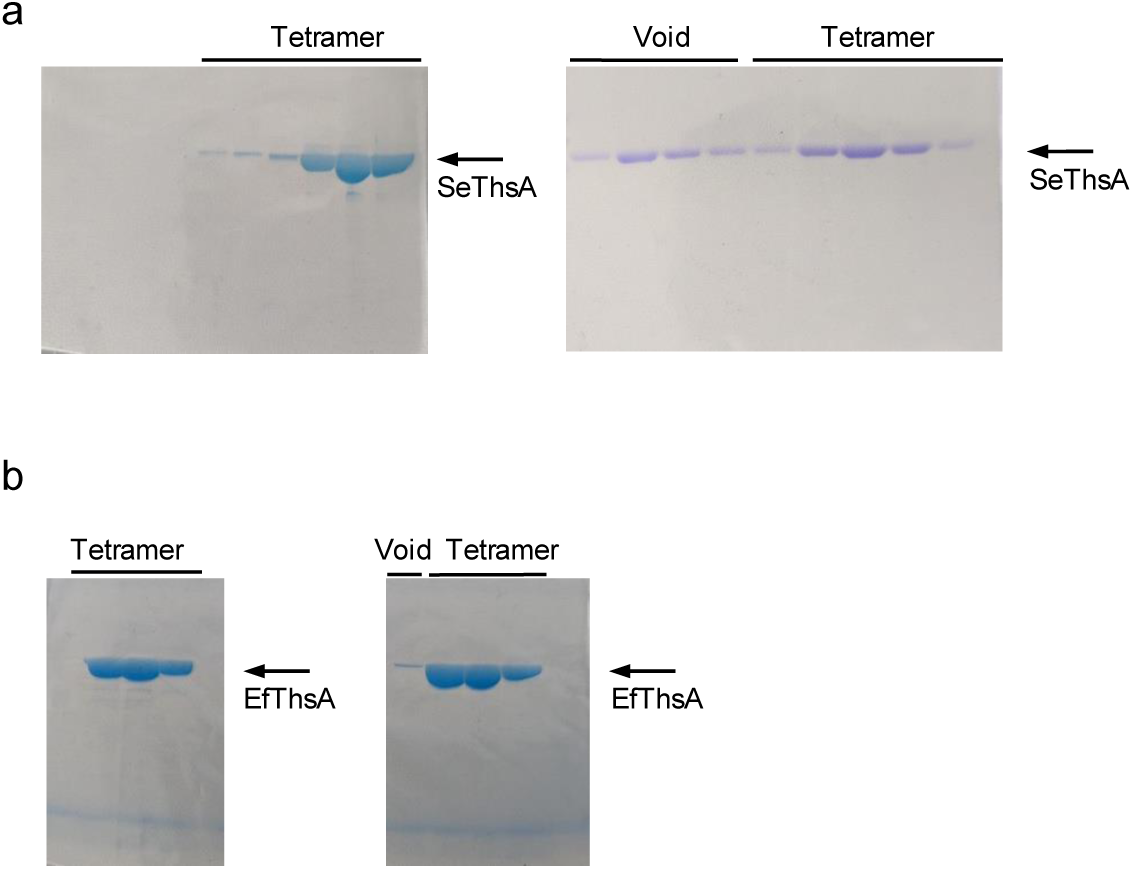
SDS-PAGE analysis of size-exclusion chromatography peaks. (a) SeThsA. (b) EfThsA. Left and right panels correspond to-/+ 3′cADPR, respectively.

**Fig. S2.**
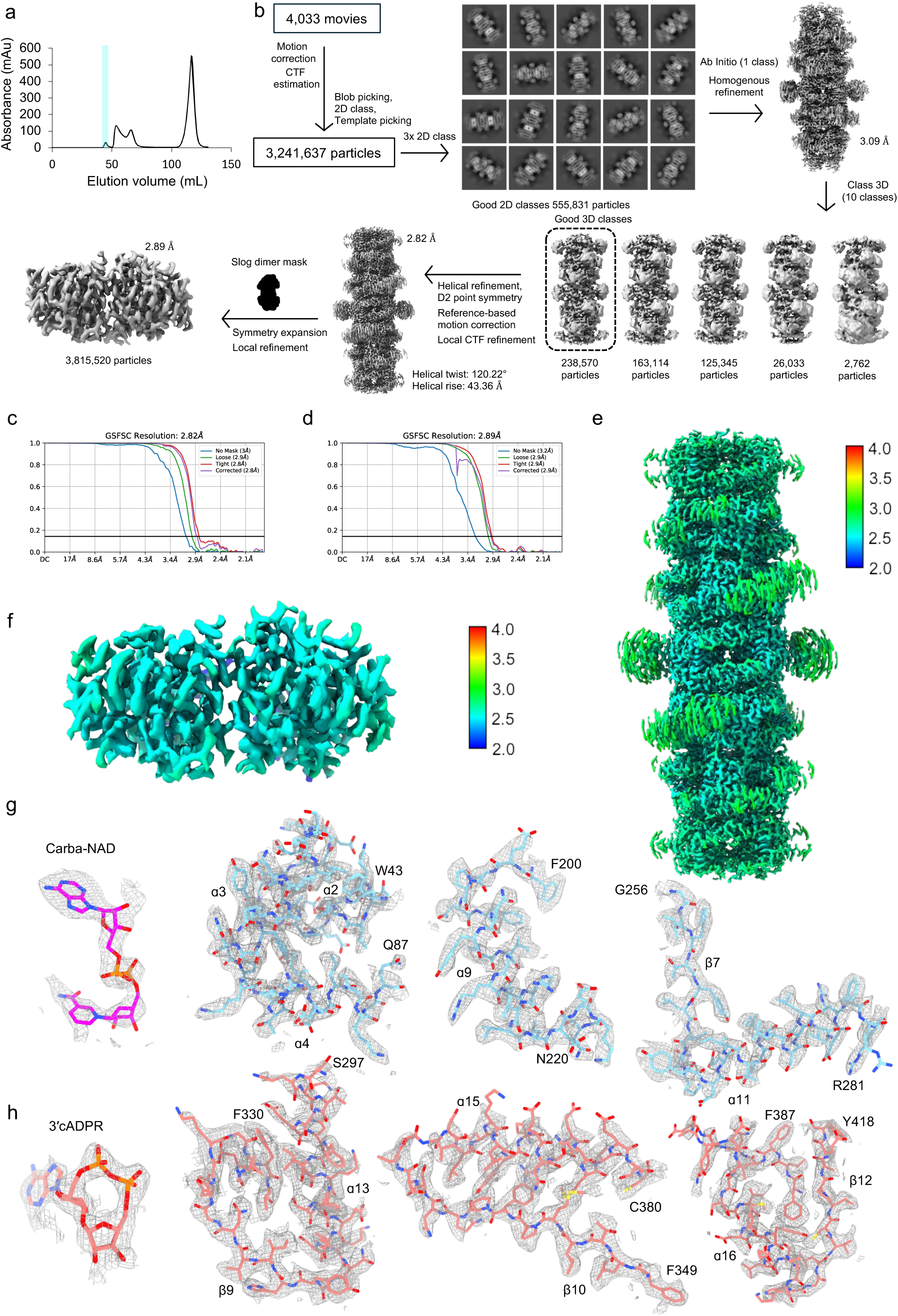
CryoEM reconstruction of SeThsA filament. (a) Size-exclusion chromatography (S200 16/60) profile of SeThsA + 3′cADPR. 80 μM SeThsA was incubated with 2 mM 3′cADPR for 1 h before size-exclusion chromatography. The fractions highlighted in blue were used for cryoEM data collection. (b) Flow-chart of the cryoEM processing steps. (c) Gold-standard FSC curves of the final 3D reconstruction of the SeThsA filament. (d) Gold-standard FSC curves of the final 3D reconstruction of the SeThsA SLOG domain dimer. (e) Local resolution map of the SeThsA filament. (f) Local resolution map of the SLOG domain dimer. (g) Representative electrostatic potential density maps for carba-NAD^+^ and the SIR2 domain of the SeThsA filament. Labelled residues indicate the N- and C-terminal residues of the displayed polypeptide. (h) Representative electrostatic potential density maps for 3′cADPR and the SLOG domain of the SeThsA filament.

**Fig. S3.**
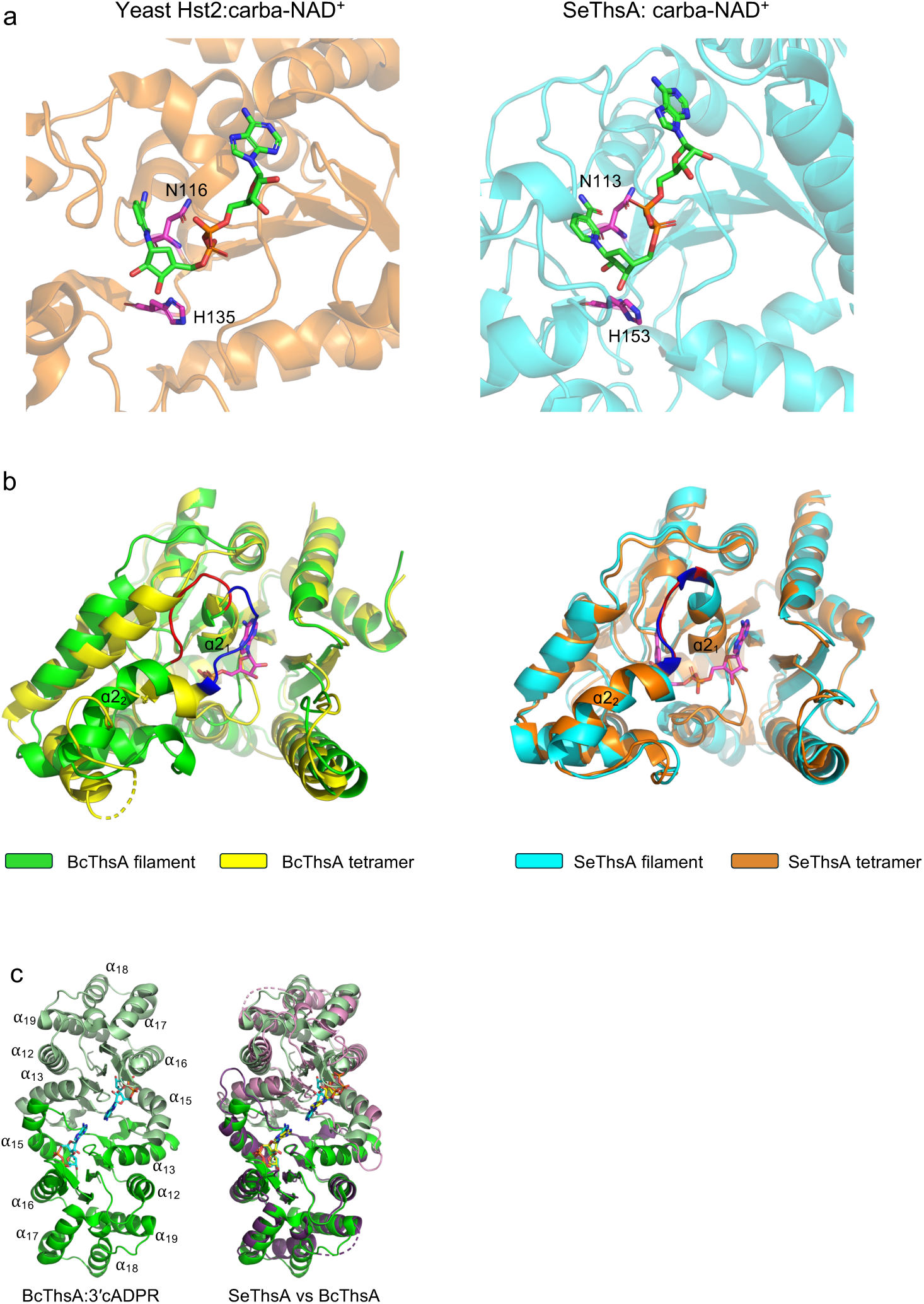
Structural analysis of SIR2 active site region and SLOG dimer interface. (a) Comparison of carba-NAD^+^ binding mode in yeast Hst2 (PDB: 1SZC) and SeThsA. (b) Comparison of the SIR2 α21-α22 loop in the tetramer and filament structures of BcThsA (PBD: 6LHX, 8BTO) and SeThsA (PDB: 7UXT). (c) Comparison of the SLOG dimers in the BcThsA (PBD: 8BTO) and SeThsA filament structures.

**Fig. S4.**
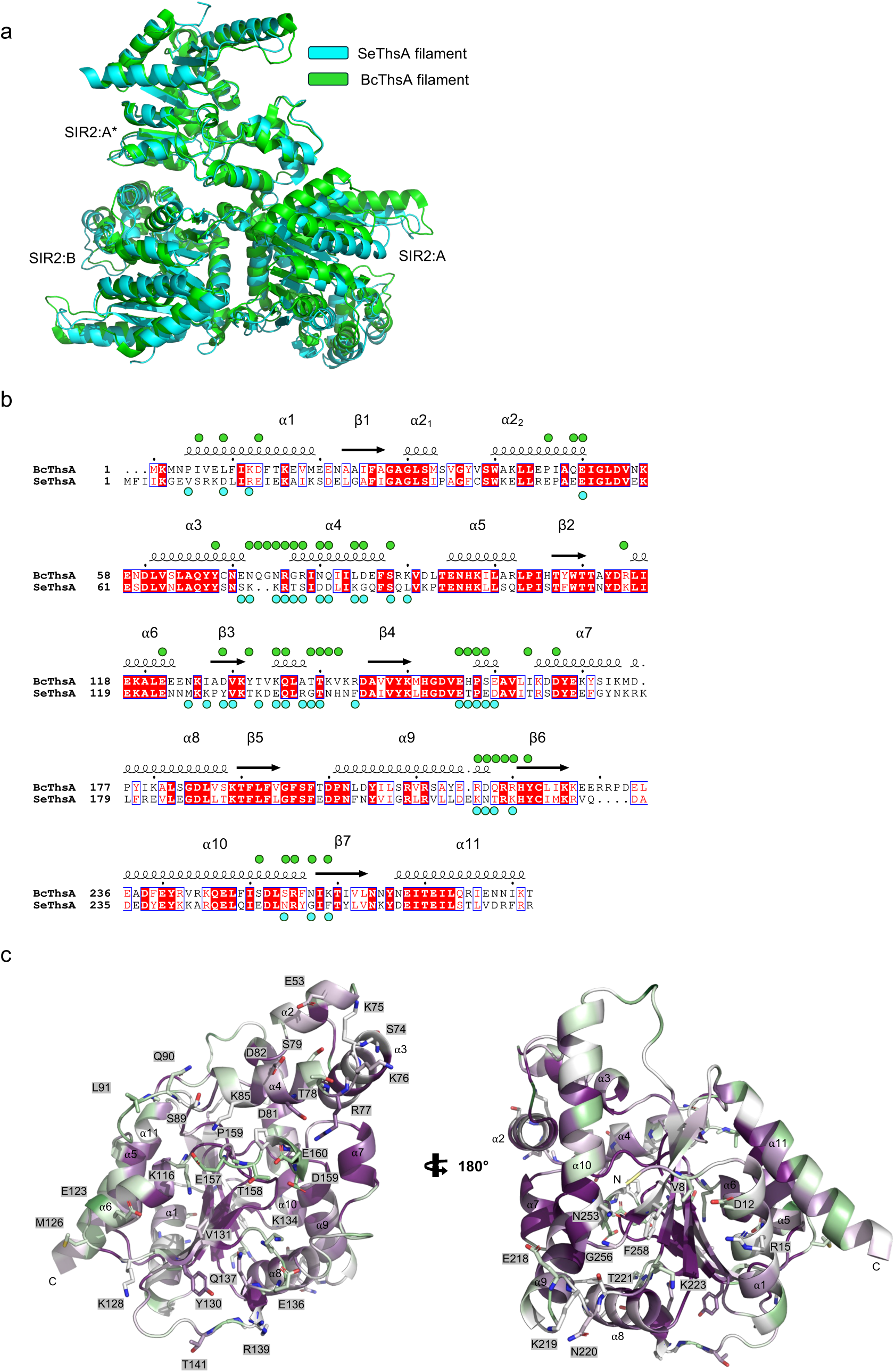
Structural analyses of the helical interface. (a) Comparison of the SeThsA (cyan) and BcThsA (green; PDB: 8BTO) helical interface. The structures were superimposed using SIR2:A*. (b) Sequence alignment of SeThsA and BcThsA. The alignment was formatted using ESPript (*55*). Strictly conserved residues are indicated in white letters with a red box and similar residues are indicated in red letters with a blue frame. Green and cyan circles indicate helical interface residues in the BcThsA and SeThsA filaments, respectively. (c) SeThsA SIR2 colored by sequence conservation using ConSurf (*56*). Green corresponds to variable regions, while purple corresponds to conserved regions. Helical interface residues are shown as sticks.

**Fig. S5.**
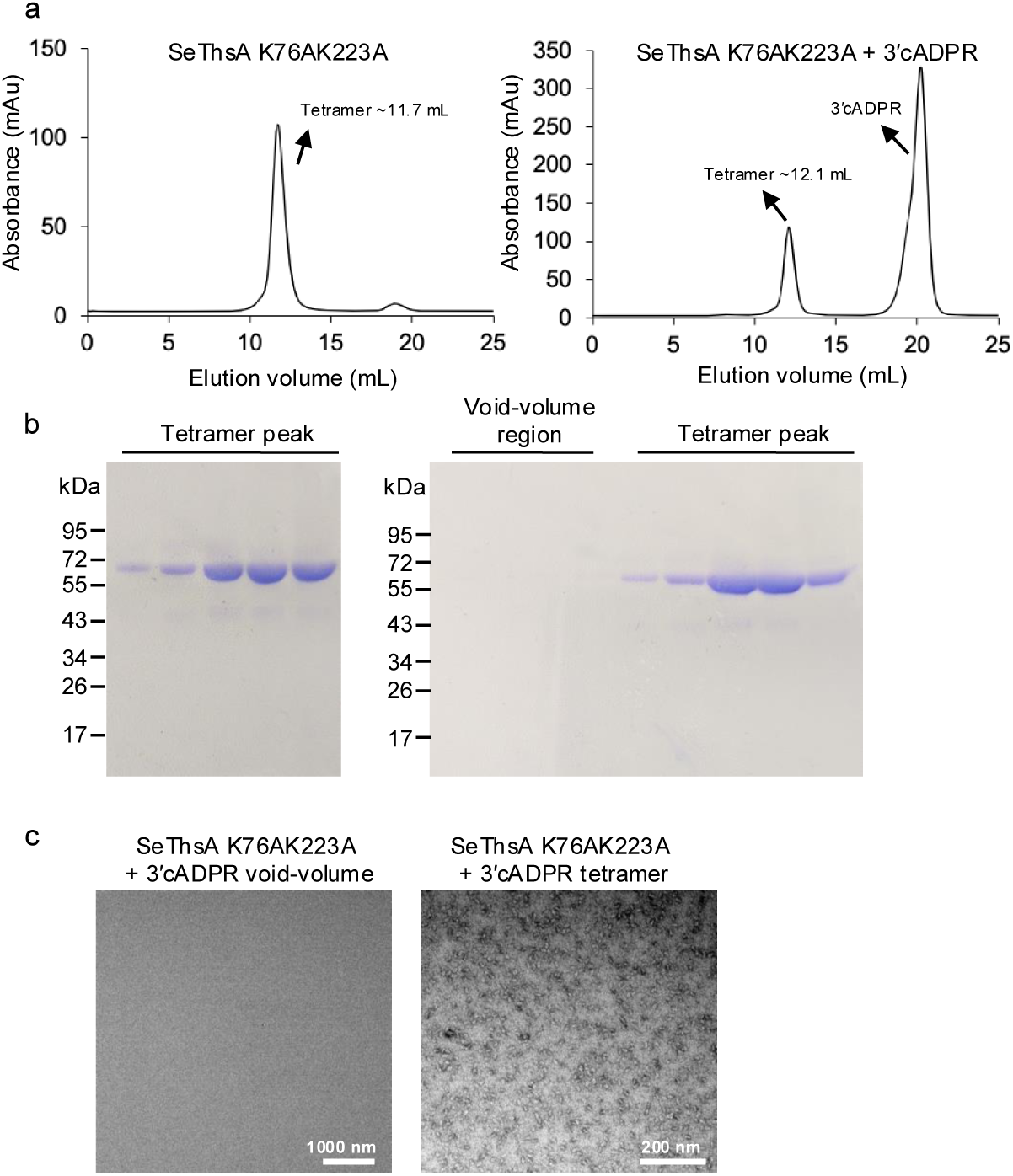
Size-exclusion chromatography and negative-stain EM analyses of the SeThsA K76AK223A mutant. (a) Size-exclusion chromatography profiles of the SeThsA K76AK223A mutant +/- 3′cADPR. (b) SDS-PAGE analyses of the tetramer peak and void-volume region in (a). Left panel: SeThsA K76AK223A; Right panel: SeThsA K76AK223A + 3′cADPR. The gels were stained with Coomassie brilliant blue. (c) Negative-stain EM images of SeThsA K76AK223A + 3′cADPR fractions. Left panel: Void volume region. Right panel: tetramer peak.

**Fig. S6.**
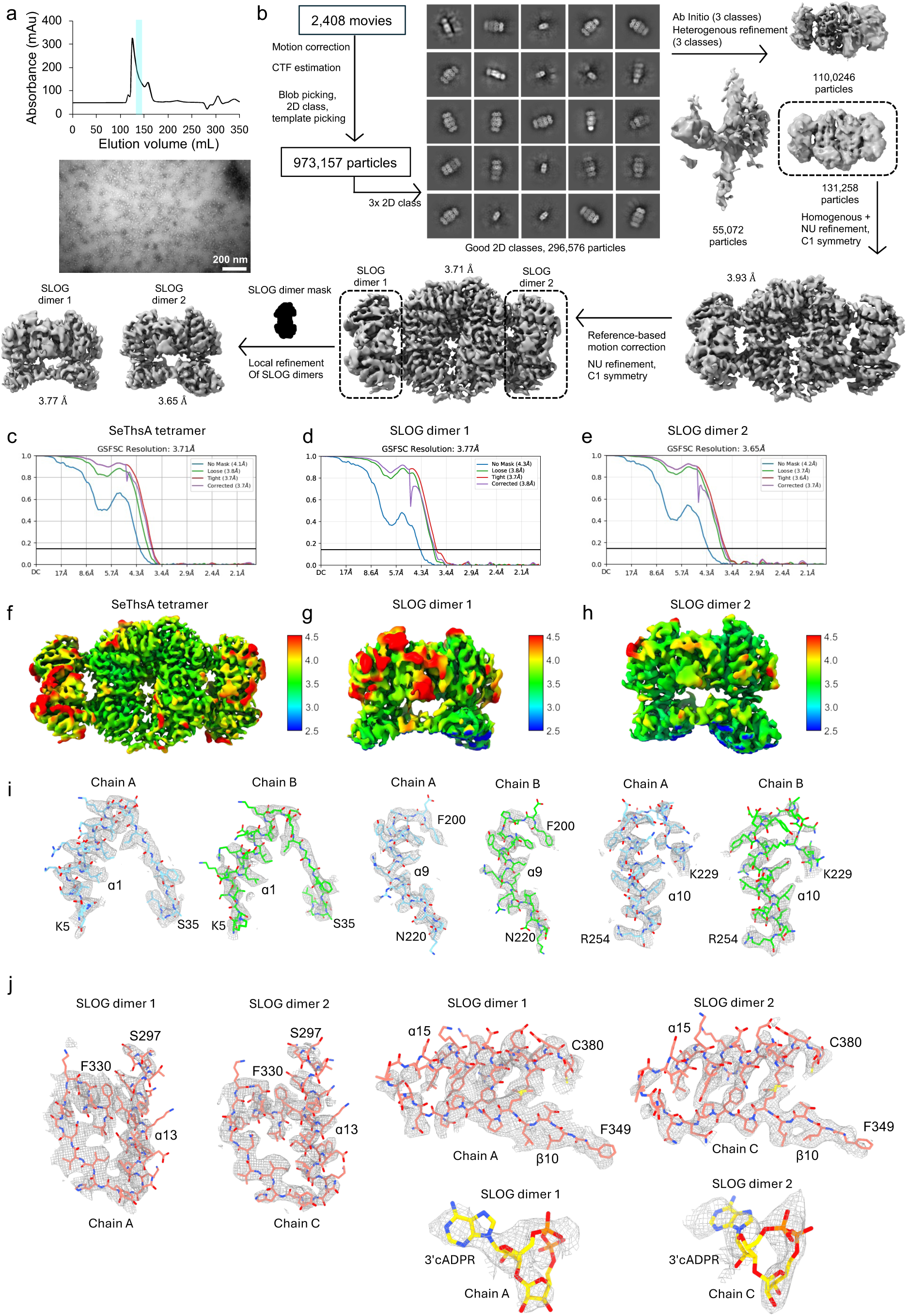
CryoEM reconstruction of SeThsA tetramer in complex with 3′cADPR. **(a) Top:** Size-exclusion chromatography (S200 26/60) profile of SeThsA + 3′cADPR. The fractions highlighted in blue were used for cryoEM data collection. 70 μM SeThsA was incubated with 1 mM 3′cADPR for 1 h before size-exclusion chromatography. Bottom: negative-stain EM analysis of the highlighted fractions. (b) Flow-chart of the cryoEM processing steps. (c) Gold-standard FSC curves of the final 3D reconstruction of the SeThsA tetramer. (d-e) Gold-standard FSC curves of the final 3D reconstruction of the SeThsA SLOG domain dimers. (f) Local resolution map of the SeThsA tetramer. (g-h) Local resolution maps of the SLOG domain dimers. (i) Representative electrostatic potential density maps for the SIR2 domain of the SeThsA tetramer. Labelled residues indicate the N- and C-terminal residues of the displayed polypeptide. (j) Representative electrostatic potential density maps for 3′cADPR and the SLOG domains of the SeThsA tetramer.

**Fig. S7.**
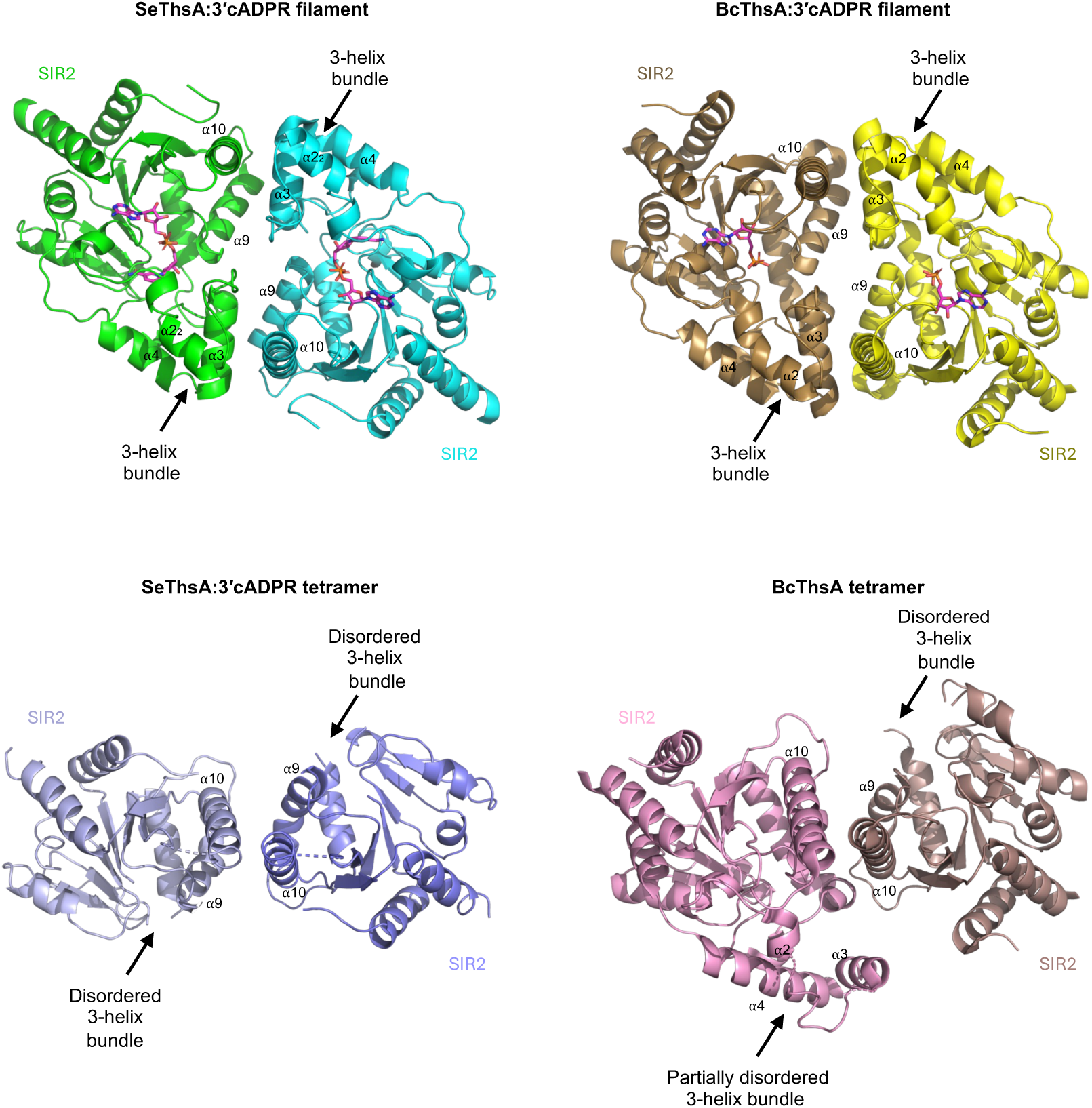
Comparison of the SIR2 dimer interfaces in the 3′cADPR-bound SeThsA tetramer cryoEM structure and the BcThsA tetramer crystal structure.

**Movie S1. Alterations to the SeThsA SLOG dimer interface upon 3′cADPR binding.** The SLOG domains are coloured in purple and pink, while 3′cADPR is displayed in yellow stick representation. The β5-α9 loops α9 helices are highlighted in yellow.

**Movie S2. Alterations to the SeThsA SIR2 tetramer interface upon 3′cADPR binding.** The SIR2 domains are coloured in green and cyan. The β5-α9 loops α9 helices are highlighted in yellow.

**Movie S3. Alterations to the SeThsA SIR2 dimer interface upon 3′cADPR binding.** The SIR2 domains are coloured in green and cyan. The β5-α9 loops α9 helices are highlighted in yellow.

---

**Table S1.** CryoEM data collection, refinement and validation statistics for SeThsA datasets.

|  | SeThsA<br>filament | SeThsA<br>filament<br>SLOG dimer | SeThsA<br>tetramer | SeThsA<br>tetramer<br>SLOG dimer 1 | SeThsA<br>tetramer<br>SLOG dimer 2 |
| --- | --- | --- | --- | --- | --- |
| **Data collection and processing** |  |  |  |  |  |
| Microscope | JEOL<br>CryoARM 300 | JEOL<br>CryoARM 300 | JEOL<br>CryoARM 300 | JEOL<br>CryoARM 300 | JEOL<br>CryoARM 300 |
| Detector | Gatan K3 | Gatan K3 | Gatan K3 | Gatan K3 | Gatan K3 |
| Voltage (kV) | 300 | 300 | 300 | 300 | 300 |
| Nominal magnification | 100,000 | 105,000 | 100,000 | 100,000 | 105,000 |
| Pixel size (Å) | 0.4864 | 0.4864 | 0.4864 | 0.4864 | 0.4864 |
| Defocus range (µm) | -0.5 to -3.0 | -0.5 to -3.0 | -0.5 to -3.0 | -0.5 to -3.0 | -0.5 to -3.0 |
| Total exposure (e/Å<sup>2</sup>) | 40 | 40 | 40 | 40 | 40 |
| Exposure per frame (e/Å<sup>2</sup>) | 1.00 | 1.00 | 1.00 | 1.00 | 1.00 |
| Total micrographs (no.) | 4,033 | 4,033 | 2,408 | 2,408 | 2,408 |
| Total extracted particles (no.) | 3,241,637 | 3,241,637 | 973,157 | 973,157 | 973,157 |
| Final particles (no.) | 238,570 | 3,815,520 | 131,258 | 131,258 | 131,258 |
| Symmetry imposed | C1 | C1 | C1 | C1 | C1 |
| Map Resolution (Å) | 2.82 | 2.89 | 3.71 | 3.77 | 3.65 |
| FSC threshold | 0.143 | 0.143 | 0.143 | 0.143 | 0.143 |
| **Refinement** |  |  |  |  |  |
| Model resolution (Å) | 2.9 | 2.9 | 3.7 |  |  |
| FSC threshold | 0.143 | 0.143 | 0.143 |  |  |
| Map correlation coefficient |  |  |  |  |  |
| Volume | 0.75 | 0.75 | 0.71 |  |  |
| Ligand (mean) | 0.63 | 0.78 | 0.46 |  |  |
| Map sharpening B-factor (Å<sup>2</sup>) | 121.9 | 134.6 | 139.3 |  |  |
| Model composition |  |  |  |  |  |
| Number of chains | 12 | 2 | 4 |  |  |
| Non-hydrogen atoms | 90960 | 5798 | 27217 |  |  |
| Residues | 5568 | 368 | 1682 |  |  |
| Water | 0 | 0 | 0 |  |  |
| Ligands | 3'cADPR: 12<br>Carba-NAD<sup>+</sup>:12 | 3'cADPR: 2 | 3'cADPR: 4 |  |  |
| B-factors (Å<sup>2</sup>) |  |  |  |  |  |
| Protein | 28.63 | 17.27 | 63.33 |  |  |
| Ligand | 35.22 | 21.04 | 93.78 |  |  |
| Bonds (RMSD) |  |  |  |  |  |
| Length (Å) (> 4σ) | 0.003 | 0.003 | 0.005 |  |  |
| Angles (°) (> 4σ) | 0.626 | 0.553 | 0.694 |  |  |
| **Validation** |  |  |  |  |  |
| Molprobity score | 0.98 | 0.98 | 1.56 |  |  |
| Clash-score | 2.10 | 2.08 | 3.99 |  |  |
| Ramachandran plot (%) |  |  |  |  |  |
| Outliers | 0.00 | 0.00 | 0.00 |  |  |
| Allowed | 1.96 | 1.11 | 5.55 |  |  |
| Favored | 98.04 | 98.89 | 94.45 |  |  |
| Rotamer outliers (%) | 0.76 | 0.63 | 0.53 |  |  |
| CB outliers (%) | 0 | 0 | 0 |  |  |
| Peptide plane (%) |  |  |  |  |  |
| Cis proline/general | 0/0/0.0 | 0/0/0.0 | 0/0/0.0 |  |  |
| Twisted proline/general | 0.0/0.0 | 0.0/0.0 | 0.0/0.0 |  |  |
| CaBLAM outliers (%) | 1.10 | 0.00 | 1.84 |  |  |

The statistics were calculated using CryoSPARC and the phenix.validation\_cryoem tool.

**Table S2.** Full nucleotide sequences of the pDR110:BcThsB, pMS039:SeThsA, pMS039:SeThsA(N113A) and pMS039:SeThsA(K76AK223A) plasmids.

